# *Ex vivo* anticoagulants affect human blood platelet biomechanics with implications for high-throughput functional mechanophenotyping

**DOI:** 10.1101/2021.02.17.428074

**Authors:** Laura Sachs, Jan Wesche, Lea Lenkeit, Andreas Greinacher, Markus Bender, Oliver Otto, Raghavendra Palankar

## Abstract

Inherited platelet disorders affecting the human platelet cytoskeleton result in increased bleeding risk. However, deciphering their impact on cytoskeleton-dependent intrinsic biomechanics of platelets remains challenging and represents an unmet need from a diagnostic and prognostic perspective. It is currently unclear whether *ex vivo* anticoagulants used for the collection of peripheral blood impact the mechanophenotype of cellular components of blood. Using unbiased, high-throughput functional mechanophenotyping of single human platelets by deformability cytometry, we found that *ex vivo* anticoagulants are a critical pre-analytical variable that differentially influences platelet deformation, their size and functional response to agonists by altering the cytoskeleton. We applied our findings to characterize the functional mechanophenotype of platelets from a patient with Myosin Heavy Chain 9 (*MYH9*) related macrothrombocytopenia. Our data suggest that platelets from *MYH9* p.E1841K mutation in humans affecting platelet non-muscle myosin heavy chain IIa (NMMHC-IIA) are biomechanically less deformable in comparison to platelets from healthy individuals.

## Introduction

Blood platelets are anucleate, discoidal multifunctional cellular fragments (1-3 µm in diameter) generated by bone marrow megakaryocytes and released into blood circulation ^1^. On exposed extracellular matrix at the sites of the vascular breach, rapid recruitment of platelets is essential for forming a primary hemostatic plug. However, under pathological procoagulatory conditions, platelets contribute to intravascular thrombosis, a leading cause of cardiovascular complications and morbidities ^2-4^. Platelets function as complex biological sensor and actuator units that respond to a broad spectrum of physicochemical stimuli via ligand-receptor-mediated interactions (i.e., outside-in signaling) and mechanotransduction events (i.e., both outside-in and inside-out signaling) ^5-7^. This complex interplay results in the coordinated regulation of signaling kinetics, including cytoskeletal remodeling that initiates platelet adhesion, activation, spreading, and platelet contraction ^8^.

It has been well established that cytoskeleton-dependent biomechanics governs diverse aspects of platelet function during hemostasis and thrombosis ^9,10^. Beyond this, the significance of platelet cytoskeletal integrity and its functional role in platelet-mediated innate immune responses such as mechano-scavenging, host defense during platelet-bacteria interactions, and vascular surveillance is emerging ^11-13^. Recent studies have also demonstrated changes in platelet biomechanical properties, and subsequent defective mechanotransduction may serve as a biophysical marker for assessing bleeding risk in individuals with inherited platelet cytoskeletal defects ^14^. Thus, deciphering cytoskeleton-dependent intrinsic biomechanical properties of platelets is highly relevant not only for broadening our understanding of the functional role of platelets in physiological and pathological processes but also from translationally significant diagnostic and prognostic perspectives ^6,10^.

Currently, a wide array of biophysical methods is available for the investigation of platelet biomechanics. They include micropipette aspiration ^15-17^, atomic force microscopy ^18-20^, scanning ion conductance microscopy ^21,22^, traction force microscopy ^23,24^, including flexible micropost arrays ^25-27^. Although these methods have proven valuable in advancing our insights into platelet biomechanics, these are technically demanding, labor-intensive, and mostly limited to analysis of adherent platelets ^28^. Besides, these methods also lack throughput, which results in implicit bias during single platelet measurements resulting from under-sampling of innate heterogeneity found in donor platelet populations ^29-31^.

The recently introduced on-chip, high-throughput real-time fluorescence and deformability cytometry (RT-FDC) has rapidly emerged as a biophysical method to address these challenges ^32,33^. RT-FDC enables continuous on-the-fly mechanophenotyping of single cells at real-time analysis rates exceeding 1000 cells/s combined with the capability of achieving molecular specificity through the application of fluorescent probes, which further opens up exciting possibilities ^34-37^.

However, on-chip deformability cytometry and other biophysical methods have not been well standardized regarding pre-analytical variability in sample preparation of cells from peripheral blood, including platelets ^38^. Specifically, it is unclear whether different *ex vivo* anticoagulants commonly used during blood sampling influence blood platelet biomechanics. Using high-throughput functional mechanophenotyping of single platelets in RT-FDC, here we demonstrate that *ex vivo* anticoagulants differentially impact intrinsic biomechanical properties (i.e., deformation and size) of human platelets. Besides this, we establish a link between platelet functional mechanophenotype, particularly their deformation and associated activation profiles as well as functional response in resting platelets and after activation with platelet agonist, respectively, in different *ex vivo* anticoagulants. We explain these findings by showing that *ex vivo* anticoagulants and platelet activation alter the content and subcellular organization of major platelet cytoskeletal components such as actin cytoskeleton and marginal band tubulin ring. Furthermore, in a potentially diagnostically significant development, using *MYH9* related macrothrombocytopenia as a model for an inherited human platelet cytoskeletal disorder, affecting platelet non-muscle myosin heavy chain IIa (NMMHC-IIA), we demonstrate that the choice of *ex vivo* anticoagulant may strongly impact the outcomes of mechanophenotyping.

## Results

### *Ex vivo* anticoagulants affect human platelet deformation and size

We first evaluated the effects of *ex vivo* anticoagulants on platelet deformation, and their corresponding size in live non-stimulated (i.e., resting) platelets in PRP by RT-FDC prepared from blood collected in ACD-A, Na-Citrate, K2-EDTA, Li-Heparin, and r-Hirudin (Fig. 1a). Non-stimulated platelets showed deformation of 0.127 ± 0.033 (mean ± SD, n=6 donors) in ACD-A, 0.111 ± 0.025 in Na-Citrate, and 0.1 ± 0.023 in r-Hirudin, which was significantly higher in comparison to the less deformable platelets at 0.071 ± 0.016 in Li-Heparin and 0.037 ± 0.01 in K_2_-EDTA. (Fig. 1b and Supplementary Fig. 4 for statistical distribution plots of individual donors). Assessment of corresponding platelet size in different *ex vivo* anticoagulants from non-stimulated platelets revealed differences in platelet size measuring at 5.035 ± 0.49 µm2 (mean ± SD, n=6 donors) in ACD-A in comparison to K2-EDTA and Li-Heparin where platelets were significantly smaller in size measuring at 4.158 ± 0.241 µm2 and 4.337 ± 0.344 µm2, respectively (Fig. 1c and Supplementary Fig. 5 for statistical distribution plots of individual donors).

**Fig 1:**
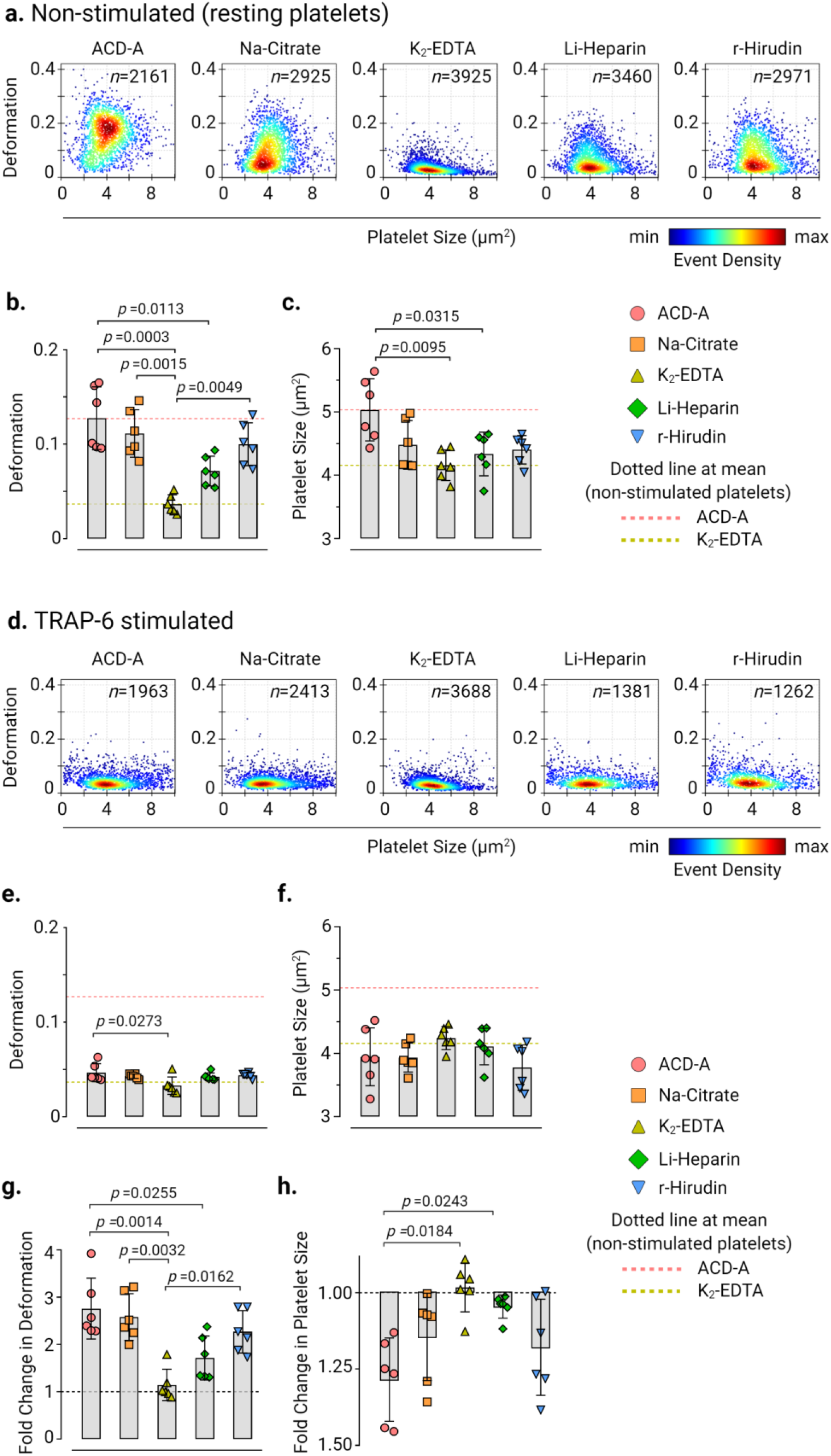
*Ex vivo* anticoagulants influence the deformability and size of human platelets. Representative KDE scatter plots of deformation and size of live single platelets in PRP prepared from whole blood collected in different *ex vivo* anticoagulants from **(a)** non-stimulated (i.e., resting) platelets and **(d)** after stimulation with TRAP-6 (*n*= number of single platelets from the same donor measured for each condition). Color coding of event density in scatter plots indicates a linear density scale from min (blue) to max (dark red). Summary data points show the median values of individual donors, and bar plots show mean ± S.D. of platelet deformation and size from non-stimulated **(b)** and **(c)** and TRAP-6 stimulated platelets **(e)** and **(f)**, respectively (n=6 donors). Fold change in platelet deformation and size upon stimulation with TRAP-6 are shown in **(g)** and **(h)**, respectively, where dotted control baseline = 1 (n= 6 donors). Statistical analysis: mixed-effects model (restricted maximum likelihood, REML) followed by Tukey’s multiple comparisons tests, with single pooled variance and *p* > 0.05 was considered significant.

Next, to test whether agonist-induced platelet activation leads to measurable changes in platelet deformation and their corresponding size depending on the type of *ex vivo* anticoagulant, TRAP-6 was used. Platelet activation by TRAP-6 resulted in a noticeable decrease in platelet deformation and a concomitant reduction in platelet size in all ex vivo anticoagulants except for K_2_-EDTA (Fig. 1d, 1e, and 1f). Assessment of fold-change in platelet deformation before and after TRAP-6 stimulation showed a decrease in platelet deformation by a factor of 2.76 ± 0.64 (mean ± SD, n=6 donors) in ACD-A, 2.58 ±0.49 in Na-Citrate, 1.72 ± 0.47 Li-Heparin, and 2.27 ± 0.45 in r-Hirudin (Fig. 1g). On the contrary, in K_2_- EDTA, TRAP-6 stimulation resulted in a minimal fold change in platelet deformation by a factor of 1.14 ±0.33 (Fig. 1g). Similarly, platelet size decreased upon TRAP-6 stimulation by a factor of 1.28 ± 0.13 (mean ± SD, n=6 donors) in ACD-A, 1.14 ± 0.14 in Na-Citrate and 1.18 ± 0.16 in r-Hirudin, while it remained unchanged at 0.98 ± 0.08 in K_2_-EDTA and 1.04 ± in Heparin (Fig. 1h). Furthermore, the changes in platelet shape observed in the RT-FDC differed between the *ex vivo* anticoagulants in non-stimulated and TRAP-6 stimulated platelets (Supplementary Fig. 6 representative bright-field images of single platelets in measurement channel overlaid with contour).

### Platelet deformation in response to platelet activation

In non-stimulated platelets, basal CD62P surface expression was not altered between all *ex vivo* anticoagulants even though platelets in K_2_-EDTA exhibited decreased deformation relative to other *ex vivo* anticoagulants (Fig. 2a, 2b, and 2c). Upon activation of platelets by TRAP-6, a significant decrease in platelet deformation with a concomitant increase in CD62P surface expression and CD62P % positive platelets was observed in all *ex vivo* anticoagulants except in K_2_-EDTA (Fig. 2d, 2e, and 2f). Assessment of fold-change in CD62P expression levels showed a significant increase by a factor of 18.19 ± 8.88 (mean ± SD, n=6 donors) in ACD-A, 21.48 ± 8.54 in Na-Citrate, 9.82 ± 7.78 in Li-Heparin, and 15.72 ±6.76 in r-Hirudin in comparison to a fold change of 2.03 ± 1.16 in K_2_-EDTA (Fig. 2g). Multivariate analysis of continuous variables from RT-FDC data from non-stimulated and TRAP-6 stimulated platelets (Fig. 2h and 2i) further confirmed platelet deformation, size, and CD62P expression is strongly affected in K_2_-EDTA.

**Fig 2:**
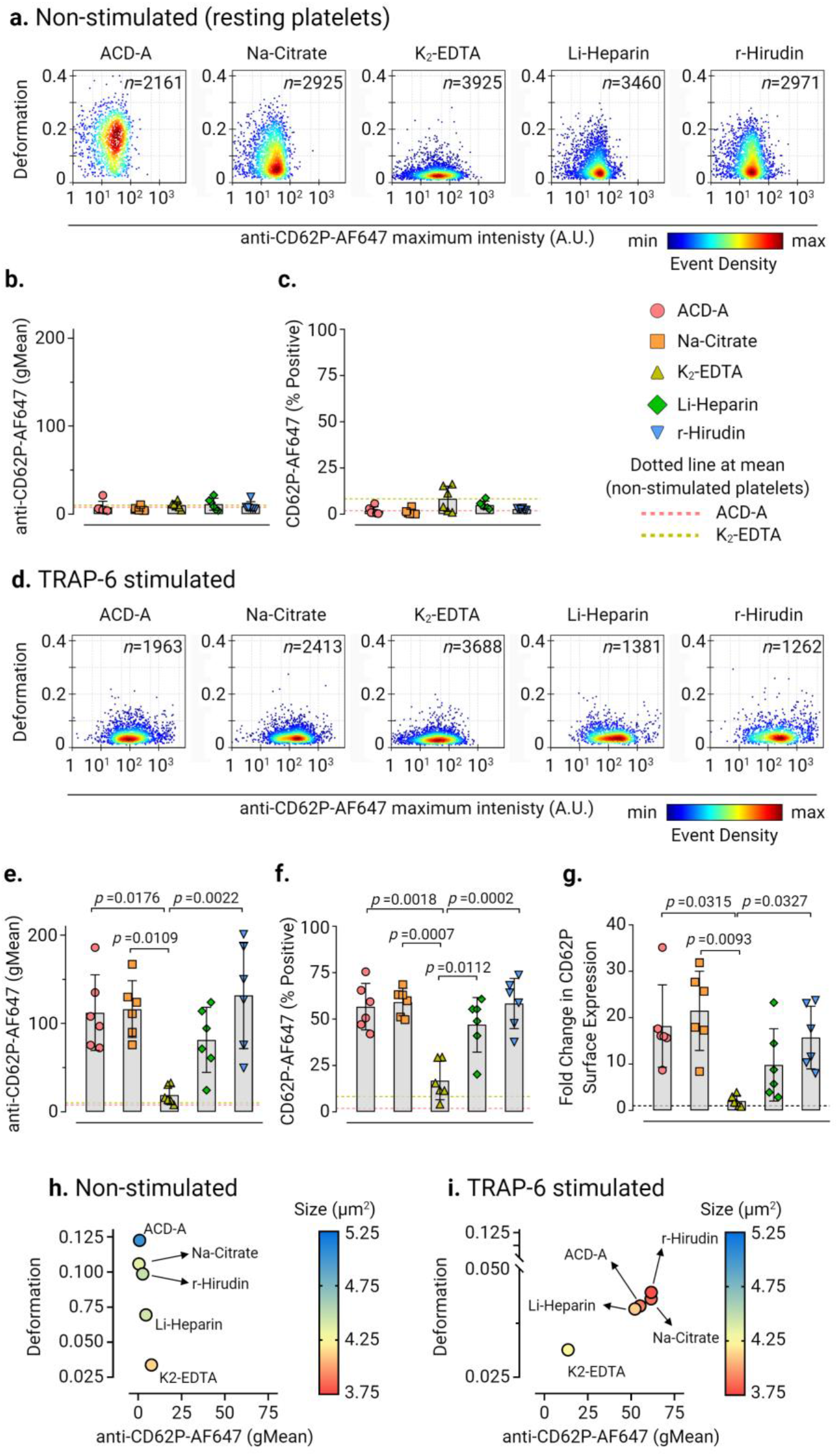
Platelet deformability and the corresponding CD62P surface expression upon activation differs in different *ex vivo* anticoagulants. Representative KDE scatter plots of platelet deformation and CD62P expression on single platelets expression (plotted on log10 scale of maximum intensity in arbitrary units (A.U.) of anti-CD62P-AlexaFluor647 antibody) in PRP different *ex vivo* anticoagulants in **(a)** non-stimulated (i.e., resting) platelets and **(d)** upon TRAP-6 stimulation. (n= number of single platelets from the same donor measured for each condition). Color coding of event density in scatter plots indicates a linear density scale from min (blue) to max (dark red). Summary graphs show of median values of individual donors, while bar graphs show mean ± S.D. of geometric mean fluorescence intensity (gMean) of CD62P expressing platelets **(b)** and **(c)** and percent positive platelets **(e)** and **(f)** above the cut-off of 5000 events or 10 min in non-stimulated and TRAP-6 stimulated platelets, respectively (n= 6 donors). Fold change in CD62P surface expression on platelets upon stimulation with TRAP-6 are shown in **(g)** where dotted control baseline = 1 (n= 6 donors). Multivariate analysis plots of continuous variables from RT-FDC, **(h)** and **(i)** of non-stimulated and TRAP-6 stimulated platelets, respectively, displaying the relationships between platelet deformability, size, and related CD62P surface expression levels in different ex vivo anticoagulants. (Data represents median values of individual variables from n= 6 donors). Statistical analysis: mixed-effects model (restricted maximum likelihood, REML) followed by Tukey’s multiple comparisons tests, with single pooled variance and p > 0.05 was considered significant.

Next, we assessed conformational changes in platelet integrin α_IIb_β_3_ as a marker for platelet activation by PAC-1 antibody binding (Fig. 3). Baseline activation levels of integrin α_IIb_β_3_ in non-stimulated platelets (Fig. 3a, 3b, and 3c) were highest in Li-Heparin (PAC-1-FITC gMean of 82.48 ± 12.16 and PAC-1-FITC % positive platelets at 36.02 % ± 9.3, mean ± S.D., n=6 donors). In contrast, K2-EDTA showed the lowest basal activation of integrin α_IIb_β_3_ of all *ex vivo* anticoagulants. In TRAP-6 stimulated platelets, PAC-1-FITC binding and PAC-1-FITC % positive platelets increased significantly (*p* < 0.0001) in all *ex vivo* anticoagulants in comparison to K_2_-EDTA (Fig. 3d, 3e, and 3f). Furthermore, fold change in PAC-1 binding to platelets after TRAP-6 stimulation increased by a factor of 2.86 ± 0.82 (mean ± SD, n=6 donors) in ACD-A, 3.39 ± 0.9 in Na-Citrate, 2.64 ± 0.7 in Li-Heparin, and 3.55 ± 0.98 in r-Hirudin in comparison to non-stimulated platelets, but did not increase in K_2_-EDTA (Fig. 3g). Also, multivariate analysis of continuous variables from RT-FDC data from non-stimulated and TRAP-6 stimulated platelets (Fig. 3h and 3i) revealed together with platelet deformation, size, and PAC-1 binding is strongly affected in K_2_-EDTA. Our results concerning the reduced binding of PAC-1 antibody to platelet integrin αIIbβ3 are consistent with previous observations in potent chelators of divalent cations such as EDTA^39^.

**Fig 3:**
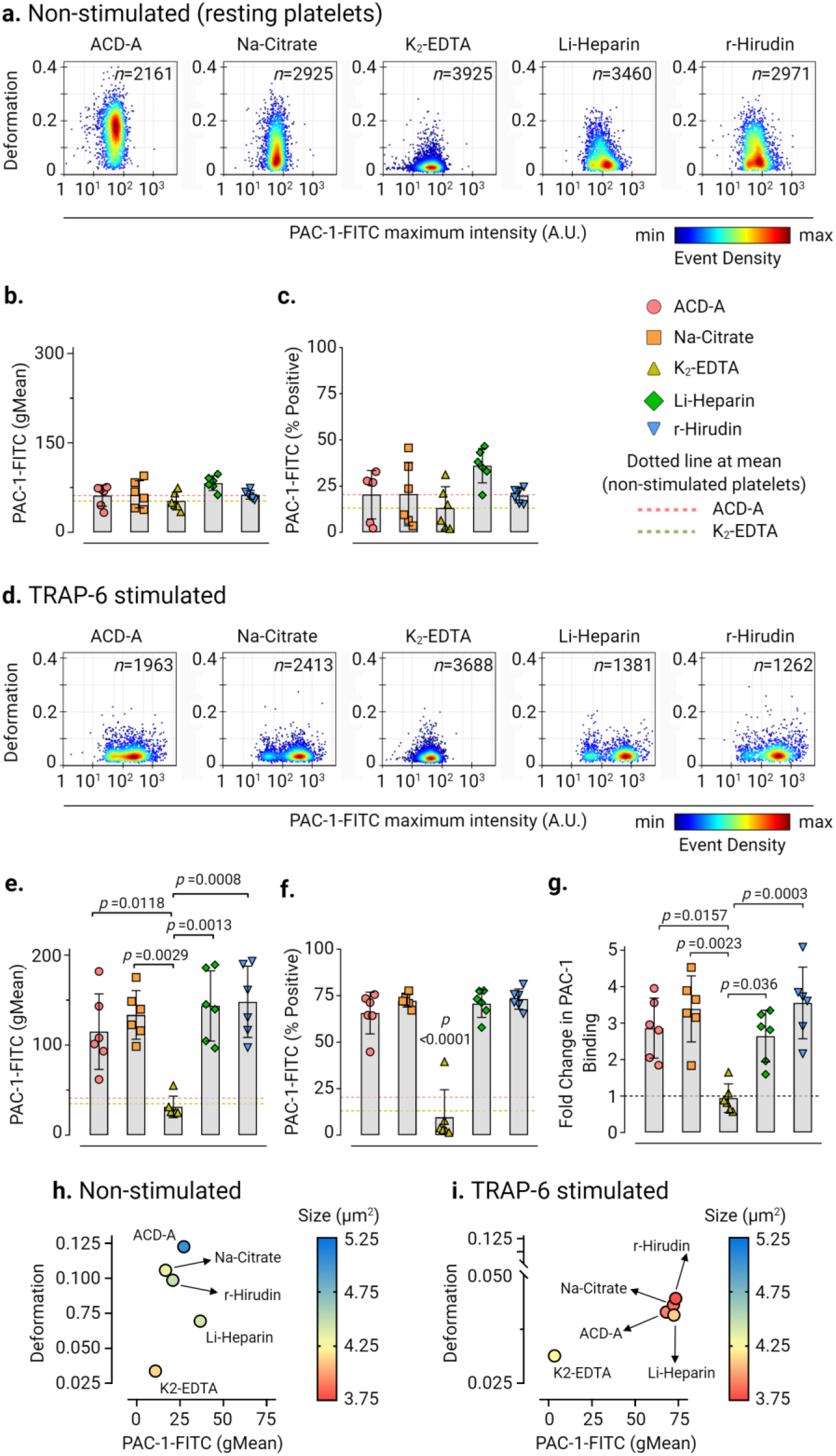
Platelet deformation and activation-induced exposure of the conformational epitope of the integrin αIIbβ3 is strongly influenced by *ex vivo* anticoagulants. Representative KDE scatter plots of platelet deformation and their corresponding activation status as a readout for binding of integrin αIIbβ3 specific ligand-mimetic PAC-1 antibody (plotted on log10 scale of maximum intensity in arbitrary units (A.U.) of PAC-1-FITC antibody) on single platelets in different *ex vivo* anticoagulants in **(a)** non-stimulated (i.e., resting) platelets and upon stimulation TRAP-6 **(b)** from a single donor (n= number of single platelets from the same donor measured for each condition). Color coding of event density in scatter plots indicates a linear density scale from min (blue) to max (dark red). Summary graphs show of median values of individual donors, while bar graphs show mean ± S.D. of geometric mean fluorescence intensity (gMean) of PAC-1-FITC antibody bound to platelets **(c)** and **(d)** and PAC-1-FITC antibody percent positive platelets **(e)** and **(f)** above the cut-off of 5000 events or 10 min in non-stimulated and TRAP-6 stimulated platelets, respectively (n= 6 donors). Fold change in CD62P surface expression on platelets upon stimulation with TRAP-6 from six donors are shown in **(g)** where dotted control baseline = 1 (n= 6 donors). Multivariate analysis plots of continuous variables from RT-FDC, **(h)** and **(i)** of non-stimulated and TRAP-6 stimulated platelets, respectively, displaying the relationships between platelet deformability, size, and PAC-1 antibody biding to integrin αIIbβ3 in different *ex vivo* anticoagulants. (Data represents median values of individual variables from n= 6 donors). Statistical analysis: mixed-effects model (restricted maximum likelihood, REML) followed by Tukey’s multiple comparisons tests, with single pooled variance and p > was considered significant.

### Increased platelet stiffness is an indicator of alterations in platelet cytoskeletal organizations and F-actin content

Fluorescence CLSM imaging and subsequent line profile analysis of non-stimulated platelets in ACD-A, Na-Citrate, Li-Heparin, and r-Hirudin showed discoidal morphology, a uniform intracellular distribution of F-actin (Phalloidin, green), and a well-defined sub-cortical marginal band microtubule ring (α-tubulin, edge-to-edge fluorescence intensity signal in magenta) (Fig. 4a and 4b). In contrast, platelets in K_2_-EDTA platelets lost their discoidal shape and were comparatively smaller. Besides, we observed an increase in subcortical localization of F-actin and coiling of microtubule ring (indicated by white arrowhead in grayscale sub-figures in Fig 4a and edge-to-edge fluorescence line profile intensity in 4b for K_2_-EDTA). Upon TRAP-6 stimulation, platelets showed substantial morphological changes compared to their non-stimulated counterparts (Fig. 4c). Line profile analysis of fluorescence intensities further revealed an increased F-actin localization at sub-cortical regions and a decrease in edge-to-edge coiling of microtubule ring in all *ex vivo* anticoagulants except for K_2_-EDTA (Fig.4d).

**Fig 4:**
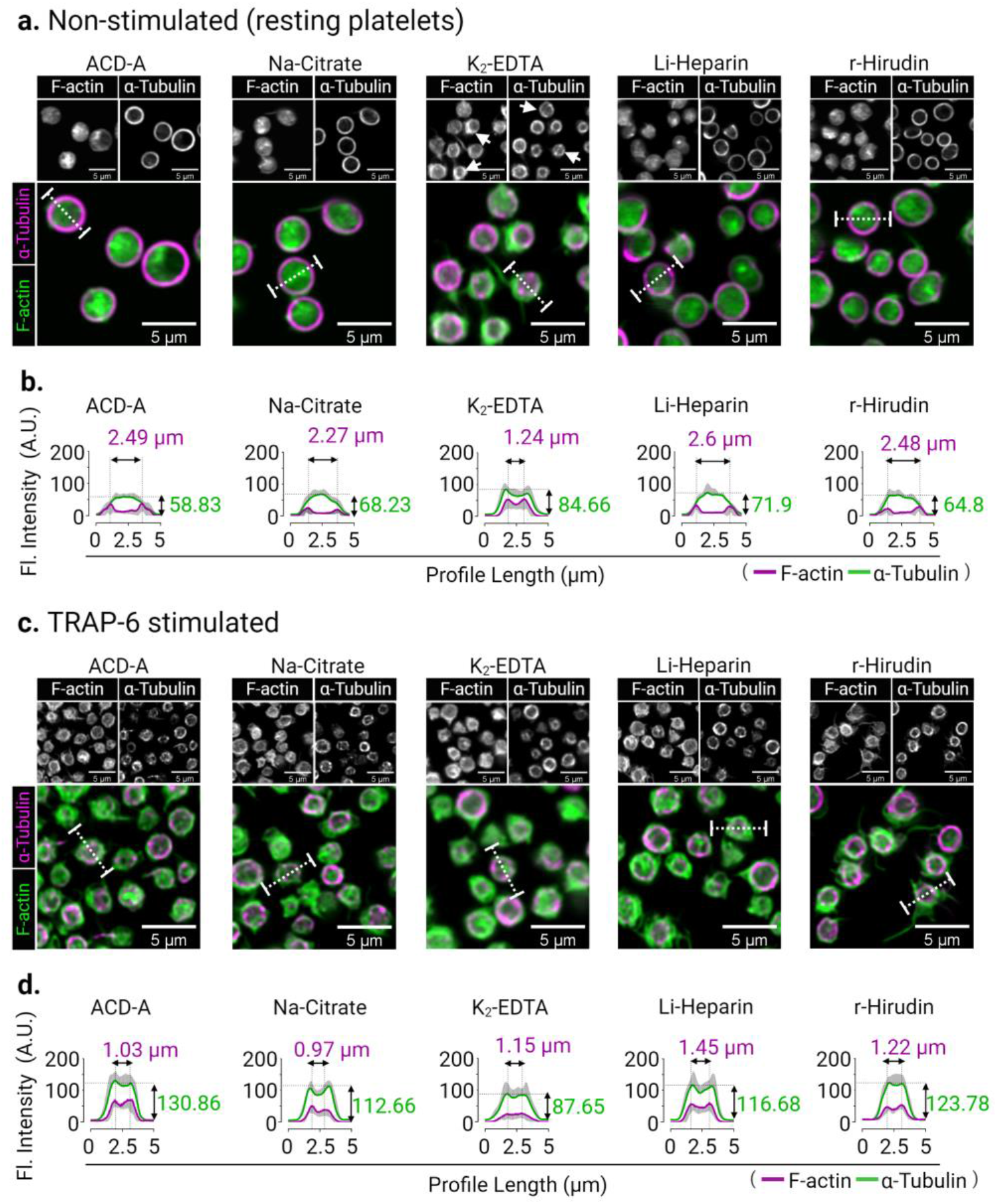
Cytoskeletal organization in resting and TRAP-6 stimulated human platelets are altered in different *ex vivo* anticoagulants. Representative confocal laser scanning fluorescence microscopic images of F-actin (green) distribution and marginal band α-tubulin (magenta) organization of human platelets in different *ex vivo* anticoagulants under **(a)** non-stimulated (resting platelets) and **(c)** 10 minutes after TRAP-6 stimulation. Fluorescence intensity line profiles (represented by the white dotted line) across individual platelets **(b)** and **(c)** show fluorescence intensity (A.U.) (along Y-axis for F-actin in green), and immunofluorescence detection of circumferential marginal band α-tubulin ring (magenta) shown as a measure of the change in the edge-to-edge length (µm) (plotted along X-axis in magenta). Graphical plots show mean ± S.D. from fluorescence intensity (A.U.) of n= 10 single platelets per *ex vivo* anticoagulant.

Next, we analyzed the total F-actin content in non-stimulated and TRAP-6 stimulated platelets in different *ex vivo* anticoagulants by flow cytometry (Fig.5a and 5b and Supplementary Fig.7). We observed a significantly higher F-actin content in non-stimulated platelets in K_2_-EDTA (Phalloidin AF647 fluorescence gMean: 162.2 ± 30.47, mean ± CD from n=6 donors) in comparison to non-stimulated platelets measuring at 110.7 ± 32.11, *p*=0.036, in ACD-A, 107 ± 23.86, *p*=0.0285 in Na-Citrate and 94.24 ±28.91, *p*=0.0023 in r-Hirudin (Fig. 5a). The basal F-actin content of platelets in Li-Heparin was found to be relatively higher (Phalloidin AF647 fluorescence gMean: 146.5 ± 9, mean ± CD from n=6 donors) than in ACD-A and Na-Citrate, statistically significant differences (*p*=0.024) were apparent between Li-Heparin and r-Hirudin in non-stimulated platelets (Fig. 5a). TRAP-6 stimulation resulted in a significant increase of total F-actin in platelets by a factor of 2.18 ± 0.28 in ACD-A, 2.13 ± 0.38 in Na-Citrate, and 2.1 ± 0.27 in r-Hirudin. In contrast, only a minor change in total F-actin content by a factor of 1.49 ± 0.26 in Li-Heparin and 0.96 ±0.14 in K_2_-EDTA was observed (Fig. 5b, 5c and Supplementary Fig.7).

**Fig 5:**
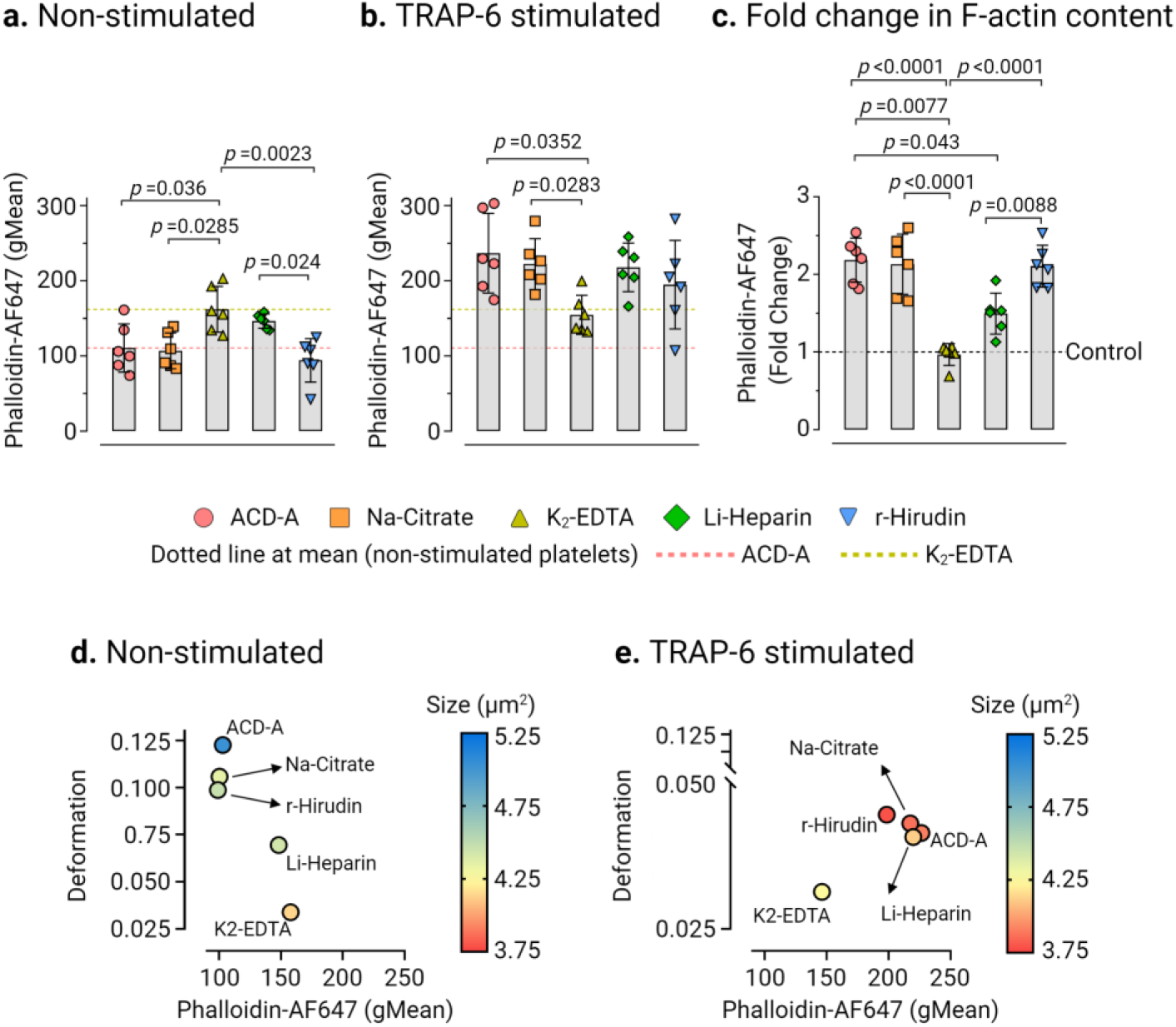
The change of F-actin content in platelets is an indicator of platelet deformability. Comparison of platelet F-actin (Phalloidin AF647 fluorescence, gMean) measured by flow cytometry in **(a)** non-stimulated and **(b)** TRAP-6 stimulated platelets in different *ex vivo* anticoagulants. Fold change in phalloidin binding **(c)** after stimulation with TRAP-6, where control baseline=1 and plots show mean ± S.D. from n=6 donors. Multivariate analysis plots of continuous variables from RT-FDC and flow cytometry, **(d)** and **(e)** of non-stimulated and TRAP-6 stimulated platelets, respectively, displaying the relationships between platelet deformability, size, and the F-actin content in different *ex vivo* anticoagulants, shown as a measure of phalloidin binding. (Data represents median values of individual variables from n= 6 donors). Statistical assessment was performed by applying the mixed-effects model (restricted maximum likelihood, REML) followed by Tukey’s multiple comparisons test, with single pooled variance and *p* > 0.05 was considered significant.

Next, a multivariate analysis of continuous variables was performed to verify whether the changes in actin polymerization status, i.e., total F-actin content measured by flow cytometry, reflect the observed differences in platelet deformation and their corresponding size by RT-FDC in different *ex vivo* anticoagulants. As shown, non-stimulated (i.e., resting) platelets (Fig. 5d) were found to deform more with low basal F-actin content in ACD-A, Na-Citrate r-Hirudin than those in Li-Heparin. Furthermore, platelets in K_2_-EDTA deformed least with higher basal F-actin content and smallest in size. Under TRAP-6 stimulation, except in K_2_- EDTA, platelets in all *ex vivo* anticoagulants showed decreased deformation, a smaller size, and increased total F-actin content (Fig 5e).

### ACD-A, but not K2-EDTA, allows mechanophenotyping of *MYH9* related disease mutations in human platelets

By RT-FDC, we next analyzed platelets from an individual with *MYH9* p.E1841K mutation in the rod region of NMMHC-IIA, an essential platelet cytoskeletal protein ^40^. In ACD-A, *MYH9* p.E1841K platelets in comparison to platelets from healthy controls deform less (0.068, median n=955 single platelets vs. 0.122, median n=2326 single platelets) and larger (5.77 µm^2^, median n=955 single platelets vs.4.05 µm^2^, median n=2326 single platelets) (Fig. 6a and Supplementary Fig.8a) under non-stimulated (i.e. resting) conditions. With TRAP-6 stimulation, platelets from the individual with *MYH9* p.E1841K in ACD-A deform further less (0.036, median n=1112 single platelets), but intriguingly their size increased (6.585 µm^2^, median n=1112 single platelets). In contrast, the platelets from the healthy individual showed decreased deformation (0.0455, median n=720 single platelets) and size (2.595µm^2^) (Fig. 6b and Supplementary Fig.8b). On the other hand, in K_2_-EDTA, non-stimulated healthy control platelets showed ≈ 3 fold decreased deformation (0.036, median n=1960 single platelets) and while *MYH9* p.E1841K platelets showed a ≥ 3 fold decreased deformation (0.0195, median n=2406 single platelets) in comparison to their counterparts in ACD-A (Fig. 6c and Supplementary Fig.9a).

**Fig 6:**
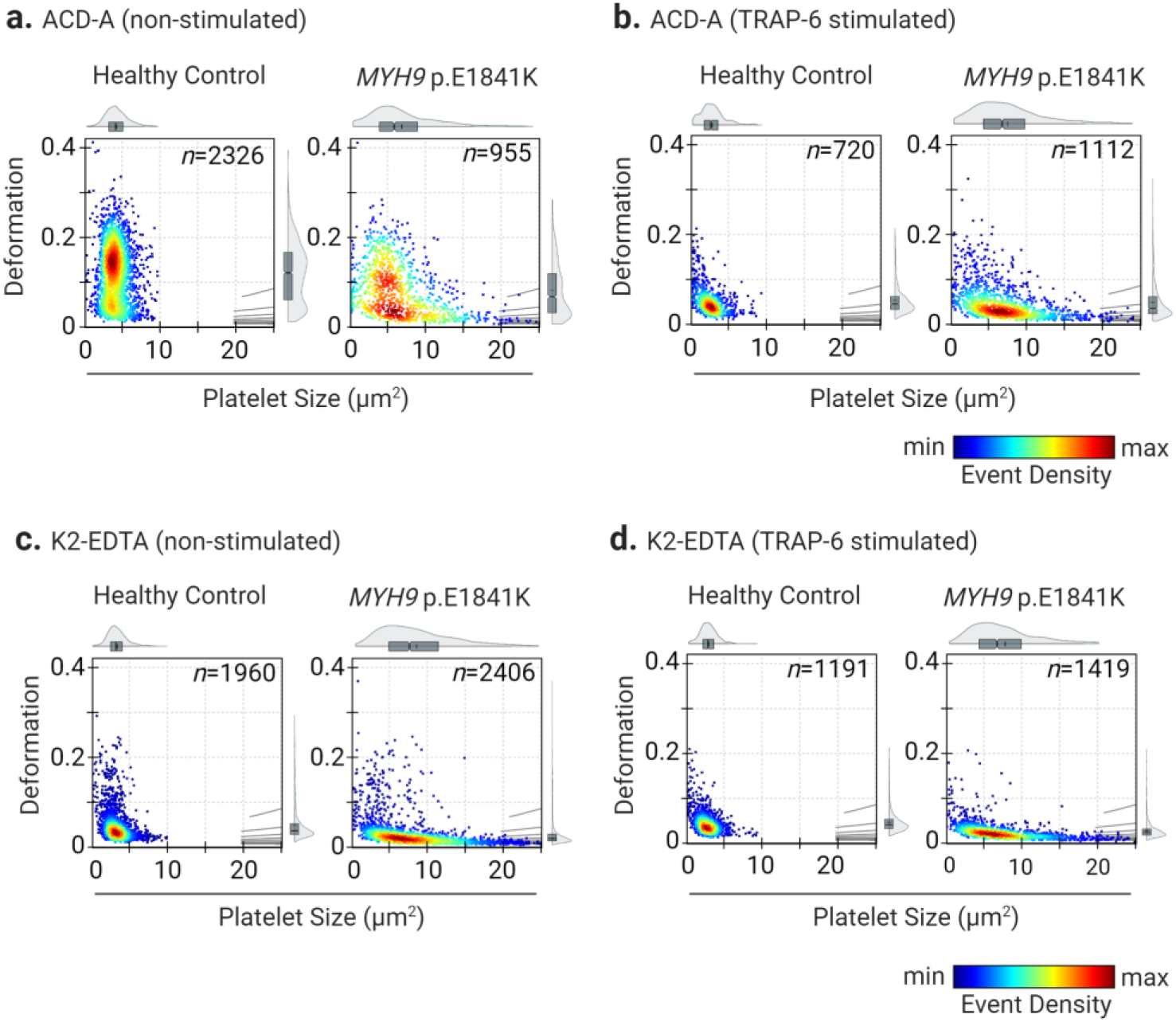
Deformability and size of platelets from a healthy individual and from a patient carrying *MYH9* p.E1841K mutation. KDE scatter plots from RT-FDC measurements performed on the same day of displaying the distribution of single platelet deformability and their corresponding size between single platelets from a healthy individual (control) and from a patient carrying the mutation *MYH9* p.E1841K for non-muscle myosin heavy chain IIa, in *ex vivo* anticoagulant **(a)** ACD-A and **(b)** in K_2_-EDTA before and after stimulation with platelet agonist TRAP-6, **(c)** and **(d)**, respectively. (n= number of single platelets). Statistical distribution plots for each condition: a notch in box plot and the horizontal line depicts median and mean, respectively, and the interquartile ranges. The full distribution of the data for each parameter is depicted by violin plots.

Besides, in K_2_-EDTA, platelets from the healthy individual became smaller (3.27µm^2^, median n=1960 single platelets). In contrast, platelets from the *MYH9* p.E1841K individual were slightly increased in their size (7.515 µm^2^, median n=2406 single platelets) compared to non-stimulated platelets in ACD-A. Furthermore, TRAP-6 stimulation of platelets in K2-EDTA only resulted in minor changes in platelet deformation and size compared to their non-stimulated counterparts (Fig. 6d and Supplementary Fig.9b). Although differences in platelet deformability and size were apparent between the platelets from *MYH9* p.E1841K patient and healthy control in ACD-A; their CD62P surface expression levels and PAC-1 binding in response to TRAP-6 were comparable.

Consistent with our observations reported above, in K2-EDTA platelets from both *MYH9* p.E1841K patient and healthy control failed to respond to TRAP-6 (Supplementary Fig.10 and Fig. 11). Furthermore, in ACD-A, assessment of F-actin content revealed a higher basal total F-actin content in non-stimulated platelets from *MYH9* p.E1841K patient at 192.86 (Phalloidin AF647 gMean) compared to the healthy control at 154.96, which upon TRAP-6 stimulation increased to 250.16 and 286.36, respectively, (Supplementary Fig.12a). On the other hand, in K_2_-EDTA, the basal total F-actin content in non-stimulated platelets from *MYH9* p.E1841K patient was found to be at 250.42 (Phalloidin AF647 gMean) and for healthy control at 206.6, that remained unchanged upon TRAP-6 stimulation (Supplementary Fig.12b).

## Discussion

The present study shows that K_2_-EDTA and Li-Heparin should not be used as *ex vivo* anticoagulants for studies on human platelet biomechanical properties. Platelets collected in ACD-A, Na-Citrate, or r-Hirudin may be used for biomechanical studies. Still, due to minor differences in the effects on platelets, results cannot be directly compared between platelets anticoagulated with these different anticoagulants. Platelets anticoagulated with Li-Heparin show some differences in their biomechanical characteristics compared to platelets in ACD-A, Na-Citrate, or r-Hirudin. Heparinized platelets show a twofold higher F-actin content, decreased deformation, and higher PAC-1 expression. The most crucial difference between Li-Heparin and r-Hirudin compared to the other anticoagulants is that Li-Heparin and r-Hirudin do not chelate calcium. However, the apparent discrepancies between Li-Heparin and r-Heparin indicate that Li-Heparin is inducing artifacts in the biomechanical properties of platelets. One explanation is the strong negative charge of heparin and its binding to αIIbβ3, which triggers αIIbβ3-mediated outside-in signals and thus initiates cytoskeletal reorganization^41,42^. Platelets collected in K_2_-EDTA have the highest F-actin content under resting conditions in comparison to the other anticoagulants and show the lowest deformation. Our observations are in agreement with previous studies, which have demonstrated K_2_-EDTA induced ultrastructural changes of the surface-bound canal system (narrowing and dilatation of the OCS) and an irreversible dissociation of the αIIbβ3 complexes^43-45 46^. It is possible that the high content of F-actin in non-stimulated and TRAP-6 stimulated platelets and the associated platelet deformation could be explained by the irreversible dissociation of the αIIbβ3 complex and the associated cytoskeletal reorganization.

The practical relevance of our findings is exemplified by the results obtained with *MYH9* p.E1841K platelets in ACD-A compared to K_2_-EDTA. The non-stimulated platelets in ACD-A deform significantly more than those in K_2_-EDTA, even after TRAP-6 induced activation. We conclude that the anticoagulants K_2_-EDTA and Li-Heparin are not suitable for the study of the human platelet cytoskeleton, while ACD-A, Na-Citrate, or r-Hirudin can be used. These results may facilitate a comparison between different laboratories using shear-based deformability cytometry such as RT-FDC to address fundamental questions of platelet physiology and its relationship with biomechanical phenotype and may help to avoid artifacts when these new technologies are applied to investigate patients with platelet disorders.

## Conclusions

In summary, we can conclude that K_2_-EDTA and Li-Heparin influence the biomechanics of platelets by decreasing the deformability and increasing actin polymerization of non-stimulated human platelets. It is recommended for the examination of the human platelet cytoskeleton to select an *ex vivo* anticoagulant such as ACD-A, Na-Citrate, or r-Hirudin and not to exchange it if possible since comparability of the results cannot be guaranteed. With the RT-FDC, we have a highly promising method to examine the platelet cytoskeleton in PRP, which according to our study, provides very solid and fast results.

## Material and Methods

### Ethics

The use of platelet-rich plasma (PRP) from healthy adult individuals and *MYH9* patients was approved by the ethics committee of the University Medicine Greifswald, Germany. All participants gave written, informed consent

### Platelet Preparation

The donors had not taken any medication in the previous ten days before blood collection. Whole blood was collected by venipuncture in BD Vacutainer^®^ Tubes containing acid citrate dextrose solution A (ACD-A), 3.8% buffered trisodium citrate (Na-Citrate), 102 I.U. Lithium-Heparin (Li-Heparin), 1.8mg/mL dipotassium ethylenediaminetetraacetic acid (K_2_ -EDTA), or 171 ATU/mL recombinant hirudin (r-Hirudin) (REVASC, Canyon Pharmaceuticals, USA). Whole blood was stored at room temperature for 15 min (at 45° angle to the horizontal surface) and then centrifuged (120 x g for 20 min at room temperature). PRP was transferred to a new polypropylene tube and incubated for 15 min at 37°C. All experimental measurements were performed within 3 hours of drawing the blood.

### Real-time fluorescence deformability cytometry (RT-FDC)

The RT-FDC setup (AcCellerator, Zellmechanik Dresden, Germany) is built around an inverted microscope (Axio Observer A1, Carl Zeiss AG, Germany) mounted with a Zeiss A-Plan 100x NA 0.8 objective. The RT-FDC fluorescence module (Supplementary Fig. 1a) is equipped with 488 nm, 561 nm, 640 nm excitation lasers, and emission is collected at the following wavelengths: 500-550 nm, 570-616 nm, 663-737 nm on avalanche photodiodes (Supplementary Fig.1a).

For functional mechanophenotyping of platelets based on molecular specificity in RT-FDC, platelets in PRP were labeled with a mouse anti-human monoclonal antibody CD61-PE (Beckman Coulter). Platelet activation was detected by direct immunofluorescence labeling of alpha granule release marker CD62P (P-selectin) with mouse anti-human monoclonal antibody CD62P-AlexaFluor647 (Clone AK4, Cat.No. 304918, BioLegend, USA). and activation associated conformational change in integrin αIIbβ3 was detected with a mouse anti-human monoclonal antibody PAC1-FITC (Clone PAC-1, Cat. No. 340507, B.D. Biosciences, USA)and, respectively. PBS (Cat.No. P04-36500, PAN Biotech GmbH, Germany) and TRAP-6 (20 µM) (Haemochrom Diagnostica GmbH, Germany) were used as vehicle control and platelet agonist, respectively. Incubations were performed at room temperature for 10 minutes in the dark.

Deformability measurements were performed in a microfluidic chip with a constriction of 15 µm x 15 μm cross-section and a length of 300 μm (Flic15, Zellmechanik Dresden, Germany) (Supplementary Fig. 1b). Platelets in suspension are injected by a syringe pump (NemeSys, Cetoni GmbH, Germany), and cell deformation occurs due to the hydrodynamic pressure gradient created by the surrounding fluid only. ^47^

Based on cellular circularity, deformation is calculated on-the-fly using bright-field images captured by a camera ^32^:

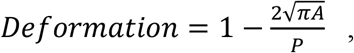

where *A* is the cross-sectional area of the cell and *P* its perimeter.

RT-FDC measurements were carried out in buffer CellCarrier B (Zellmechanik Dresden, Germany), which is composed of 0.6% (w/v) methylcellulose in PBS (without Ca^2+^, Mg^2+^). Here, 50 μL of immunofluorescently labeled PRP was suspended in 450 μL CellCarrier B. The PRP suspension was then driven through the microfluidic chip at flow rates of 0.006 μl/s, and the measurement was stopped after achieving 5000 single platelet count (hard-gate 150-33000 arbitrary units, A.U. for CD61-PE of fluorescence intensity) or after 10 min. RT-FDC data was acquired using the ShapeIn software (Version 2.0, Zellmechanik Dresden, Germany). Using the Shape-Out analysis software (https://github.com/ZELLMECHANIK-DRESDEN/ShapeOut2/releases/tag/2.3.0 Version 2.3, Zellmechanik Dresden, Germany), kernel density estimation (KDE) plots of event density were generated, and statistical analysis was performed to determine the median values for platelet deformation, their size and the geometric mean of fluorescence (gMean) of the relevant functional variables. The range area ratio was limited to 0 - 1.1 and the cell size to 0-10 μm for the analysis (Supplementary Fig. 2 and 3).

### Flow cytometry

Platelets were treated as described above for RT-DC. We used PerFix-nc Kit (Cat.No. B31167; Beckman Coulter GmbH, Germany) and Phalloidin-Atto-647 (Atto-Tec GmbH, Germany) to measure changes in total F-actin content in the platelets. Flow cytometry data were processed using FlowJo™ software for Windows, Version v10.6.2. (Becton, Dickinson and Company, USA), and the gMean of the relevant variables was determined.

### Fluorescence Microscopy

Platelets in PRP were incubated with PBS (vehicle control, non-stimulated)) or stimulated with TRAP-6 for 10min followed by fixation in 2% paraformaldehyde (Morphisto, Germany) for 15min. Fixed platelets were transferred into a Shandon™ Single Cytofunnel™ (Thermo Fisher, USA) and were centrifuged on a microscope slide for 5 min at 700rpm (Cytospin ROTOFIX 32 A, Hettich, Germany), washed thrice with PBS (5 min intervals). Platelet were permeabilized in 0.5 % saponin with 0.2% bovine serum albumin (BSA) (Cat. No. 11924.03, SERVA Electrophoresis GmbH, Germany) for 25 min, and followed by blocking for 30 min in 0.5% saponin supplemented with heat-inactivated 10% normal goat serum. Permeabilized platelets were incubated with 1:500 dilution of mouse monoclonal anti-α-Tubulin IgG (Clone DM1A, Cat.No. T9026, Sigma Aldrich GmbH, Germany) primary antibody diluted in 0.5% saponin with 0.2% BSA in PBS for 16 hours at 4°C followed by three washing steps in PBS for 5 min each. Afterward, platelets were incubated with 1:750 dilution of goat polyclonal anti-mouse AF488 IgG prepared in 0.5% saponin with 0.2% BSA in PBS for 60 minutes in the dark at room temperature, followed by three washing steps with PBS for 5 min each. F-actin was labeled with 20 pM Phalloidin Atto 647N (Cat. No. AD 647N-81, Atto-Tec GmbH, Germany) for 60 minutes, followed by three washes in PBS for 5 minutes each. Slides were covered by a permanent mounting medium Roti®-Mount FluorCare (Cat. No. HP19.1, Karl-Roth GmbH, Germany). Fluorescence microscopy was performed on a Leica SP5 confocal laser scanning microscope (Leica Microsystems, Wetzlar, Germany) equipped with HCX PL APO lambda blue 40x/1.25 OIL UV objective. For image acquisition, AF488 and ATTO647 were excited by argon (488nm) and helium-neon (HeNe) (633nm) laser lines selected with an acousto-optic tunable filter (AOTF), and fluorescence emission was collected between 505-515 nm and 640-655 nm respectively on hybrid detectors (HyD). Assessment of F-actin distribution and organization of marginal band α-tubulin staining was performed by measuring cross-sectional line profile (5 µm length and 1 µm width) of non-saturated grayscale fluorescence intensities (pixel values) of immunofluorescent probes across individual platelets in confocal images using Leica Application Suite X (Version 3.7.1, Leica Microsystems, Wetzlar, Germany). For data plotting, GraphPad Prism version 8.0.0 for Windows (GraphPad Software, San Diego, California USA) was used.

### Statistical plots and analysis of RT-FDC data

Statistical plots showing parameters of the platelet population were prepared with ShapeOut software (https://github.com/ZELLMECHANIK-DRESDEN/ShapeOut2/releases/tag/2.3.0 Version 2.3, Zellmechanik Dresden), PlotsOfDifferences (https://huygens.science.uva.nl/PlotsOfDifferences/) and Raincloud Plots (https://gabrifc.shinyapps.io/raincloudplots/). ^48,49^ Statistical assessment was performed using the mixed-effects model (restricted maximum likelihood, REML) followed by Tukey’s multiple comparisons test, with single pooled variance using GraphPad Prism version 8.0.0 for Windows (GraphPad Software, San Diego, California USA). *P*<0.05 was considered significant.

## Supporting information

Supplementary Fig. 1. Schematics of RT-FDC setup.

## Data availability

Source data and accompanying software for analysis of RT-FDC data are available in a citable repository and can be accessed by requesting the corresponding author(s).

## Acknowledgments

O.O gratefully acknowledges funding from the German Ministry of Education and Research (BMBF) within the project 03Z22CN11 (ZIK grant) and the German Center for Cardiovascular Research within the project 81×3400107 (Postdoc start-up grant). This work was supported by the Deutsche Forschungsgemeinschaft project number 374031971–CRC/TR 240 project A06 to M.B., O.O. and R.P.

## Author contributions

L.S., A.G.,M.B.,O.O. and R.P. designed the study. L.S. performed all RT-FDC experiments. J.W. and L.L. performed flow cytometry. L.S. and R.P. performed CLSM experiments. L.S., and R.P. analyzed the data and prepared figures. L.S. wrote the manuscript. A.G. provided access to *MYH9* patient platelets. A.G., M.B., O.O. and R.P. contributed to writing the manuscript. All authors contributed to the critical revision of the manuscript. .M.B., O.O. and R.P. obtained funding.

