## Supplementary Fig. 1. Schematics of RT-FDC setup. for "*Ex vivo* anticoagulants affect human blood platelet biomechanics with implications for high-throughput functional mechanophenotyping"

*corresponding authors

**Contents:**

**Supplementary Fig. 1. Schematics of RT-FDC setup.**

**Supplementary Fig. 2: Calculation of Area Ratio *Δ* in Shapeout software.**

**Supplementary Fig. 3: Application of range area ratio and cell size filter to raw data acquired in RT-FDC by using in ShapeOut software.**

**Supplementary Fig. 4: Distribution of single platelet deformation in individual donors.**

**Supplementary Fig. 5: Distribution of single platelet size in individual donors.**

**Supplementary Fig. 6: Event images of platelet morphologies in the RT-FDC measurement channel.**

**Supplementary Fig. 7: Flow cytometric detection of platelet CD62P expression and F-actin content.**

**Supplementary Fig.8: Event images with the contour of platelet morphologies from the RT-FDC measurement channel from healthy individual and MYH9 p.E1841K patient in ACD-A.**

**Supplementary Fig. 9: Event images with the contour of platelet morphologies from the RT-FDC measurement channel from healthy individual and MYH9 p.E1841K patient in ACD-A. in $K_2$-EDTA.**

**Supplementary Fig.10: Platelet deformation and CD62P expression from a patient with *MYH9* p.E1841K.**

**Supplementary Fig. 11: Platelet deformation and platelet activation measured for binding of the anti-PAC-1 antibody as a function of exposure of conformational epitope of the integrin αIIbβ3 from a patient with *MYH9* p.E1841K.**

**Supplementary Fig.12: Flow cytometric detection of platelet CD62P expression and F-actin content in *MYH9* p.E1841K patient.**

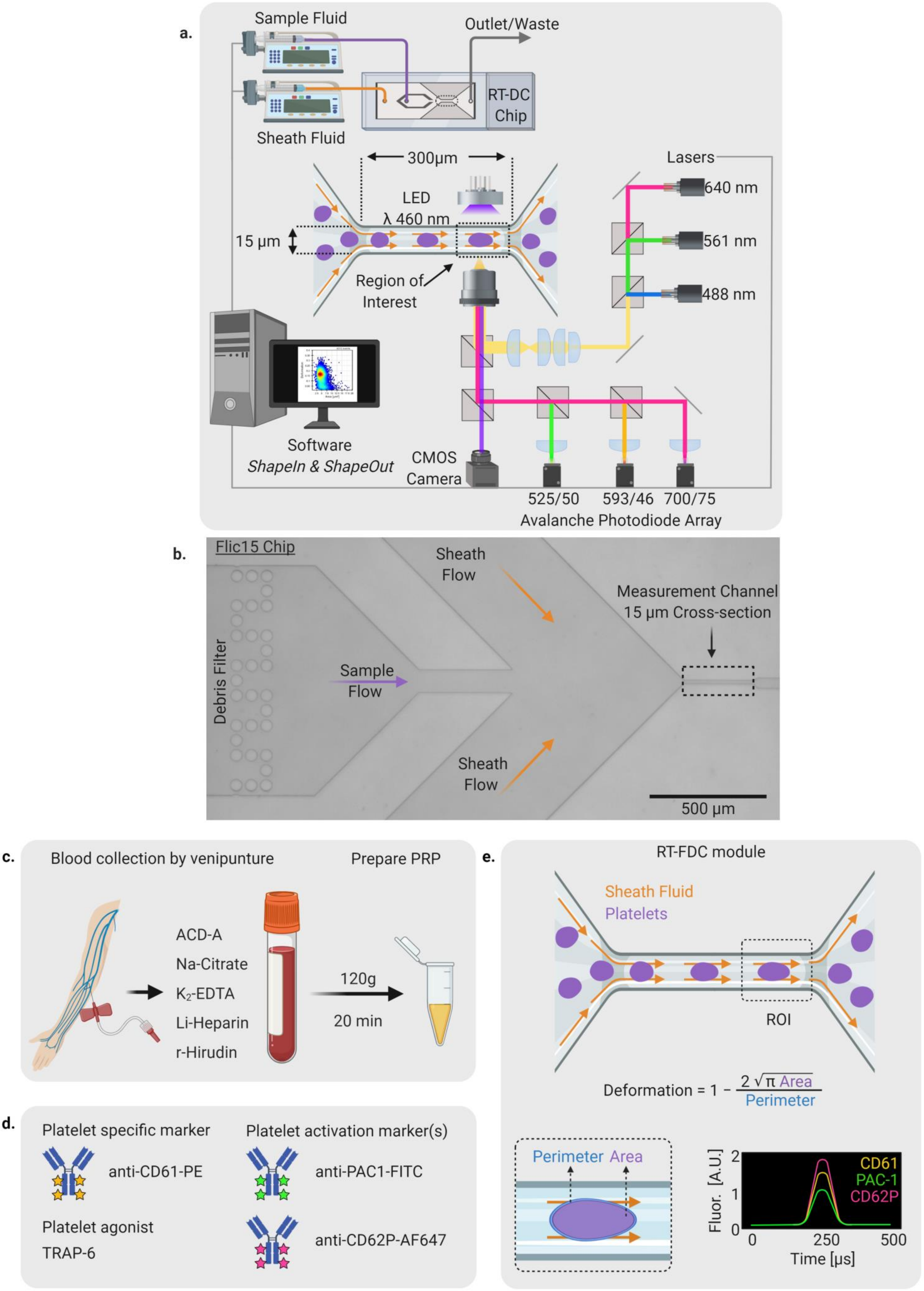

a.
Sample Fluid
Outlet/Waste
RT-DC Chip
Sheath Fluid
300µm
LED
λ 460 nm
15 µm
Region of Interest
Lasers
640 nm
561 nm
488 nm
Software
ShapeIn & ShapeOut
CMOS Camera
525/50
593/46
700/75
Avalanche Photodiode Array
b.
Flic15 Chip
Sheath Flow
Measurement Channel
15 µm Cross-section
Debris Filter
Sample Flow
Sheath Flow
500 µm
c.
Blood collection by venipunture
Prepare PRP
ACD-A
Na-Citrate
K2-EDTA
Li-Heparin
r-Hirudin
120g
20 min
d.
Platelet specific marker
Platelet activation marker(s)
anti-CD61-PE
anti-PAC1-FITC
Platelet agonist
TRAP-6
anti-CD62P-AF647
e.
RT-FDC module
Sheath Fluid
Platelets
ROI
Deformation = 1 − 2 √π Area / Perimeter
Perimeter
Area
Fluor. [A.U.]
CD61
PAC-1
CD62P
Time [µs]

**Supplementary Fig. 1: a.** Schematic depiction of real-time fluorescence and deformability (RT-FDC) instrumentation configuration and **b.** overview of the Flic15 microfluidic chip used for mechanophenotyping of human platelets showing the arrangement of micro-channels for the dedicated flow of sample and sheath fluids that merge into the measurement channel with a cross-section of 15 µm and a length of 300 µm. Graphical description of the experimental outline for functional mechanophenotyping of single platelets by RT-FDC.. **c.** Sample Preparation: Whole blood is collected by venipuncture into *ex vivo* anticoagulants: ACD-A, sodium citrate, $K_2EDTA$, Li-Heparin, and r-Hirudin. Freshly prepared platelet-rich plasma (PRP) from whole blood by centrifugation is mixed with **d**. antibodies and with/without agonists. c Platelet suspension is pumped through a microfluidic channel and imaged by CMOS camera, while three lasers excite and avalanche photodiodes measure fluorescence. In the narrow constriction zone, cell deformation occurs due to the hydrodynamic pressure gradient created by the surrounding fluid deformation is determined by on-the-fly image processing.

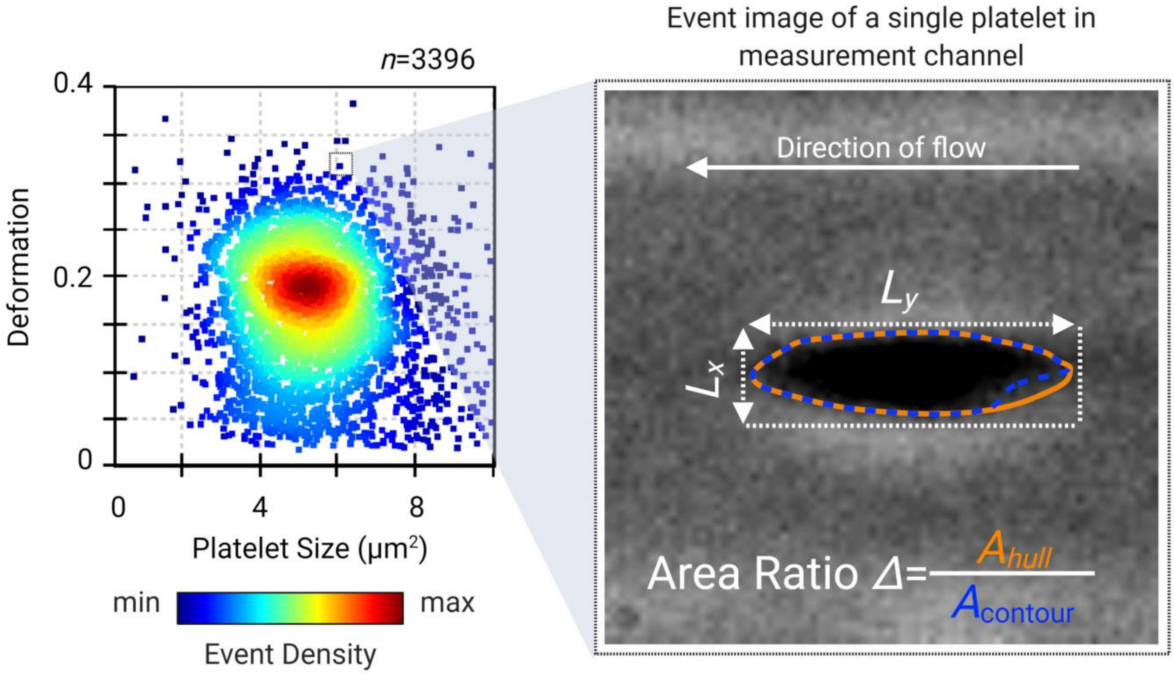


**Supplementary Fig. 2: Calculation of Area Ratio *Δ* in Shapeout software.** Example RT-FDC scatter plot of platelet deformation against size from non-stimulated platelets, colored by Kernel Density Estimate (KDE), showing a single platelet's bright-field event image during passage through microfluidic measurement channel of 15µm in cross-section. The bounding box defines the lengths *Lx* and *Ly*, and the area of the contour ($A_{contour}$) and convex hull ($A_{hull}$) are used to calculate the platelet size and circularity, respectively. The ratio between $A_{hull}$ and $A_{contour}$ gives the area ratio (*Δ*), used for applying the filter to select only platelets for further analysis.

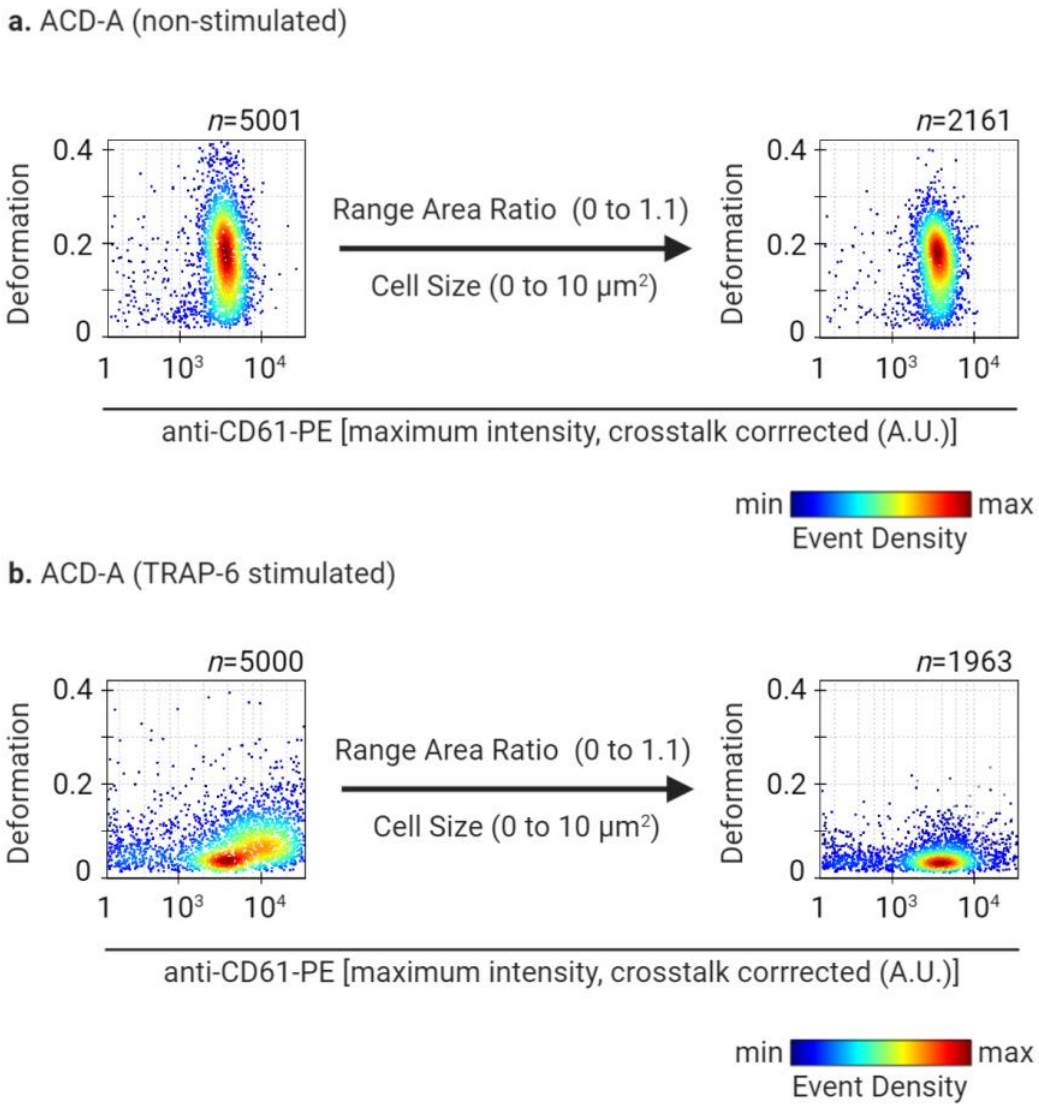


**Supplementary Fig. 3: Application of range area ratio and cell size filter to raw data acquired in RT-FDC by using in ShapeOut software.** Scatter Plot colored by Kernel Density Estimate (KDE) of RT-FDC data plotted for platelet deformation (along Y-axis) and for fluorescence intensity of platelets specifically labeled for CD61 surface with mouse monoclonal anti-human CD61-PE marker (along X-axis, maximum fluorescence intensity, crosstalk corrected (A.U.)) before and after applying filters for range area ratio and cell size in ShapeOut software in non-stimulated (a) and TRAP-6 stimulated platelets (b).

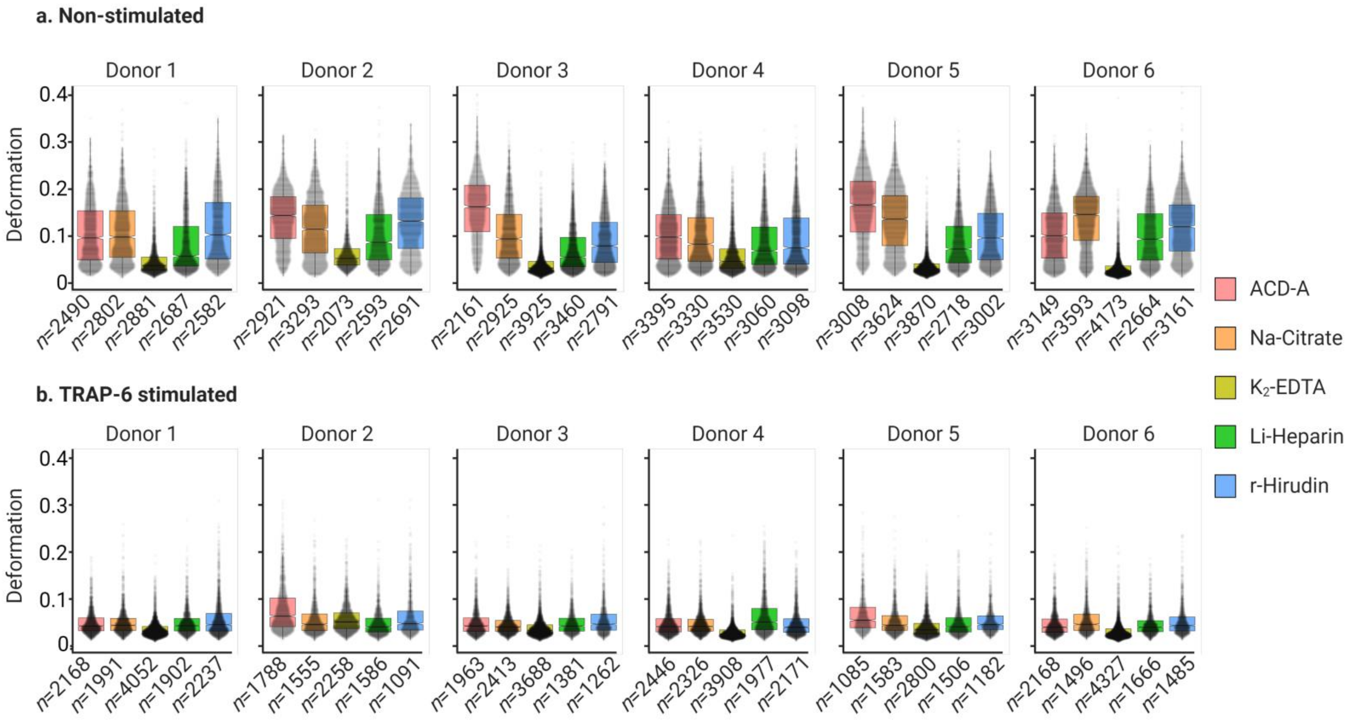


**Supplementary Fig. 4: Distribution of single platelet deformation in individual donors.** Violin plots of platelet deformation from six healthy donors before and after stimulation with PAR-1 agonist TRAP-6 peptide (20µM) for 10 minutes in the presence of PRP obtained from blood collected in different anticoagulants. n= number of single platelets analyzed for each per donor for each *ex vivo* anticoagulant.

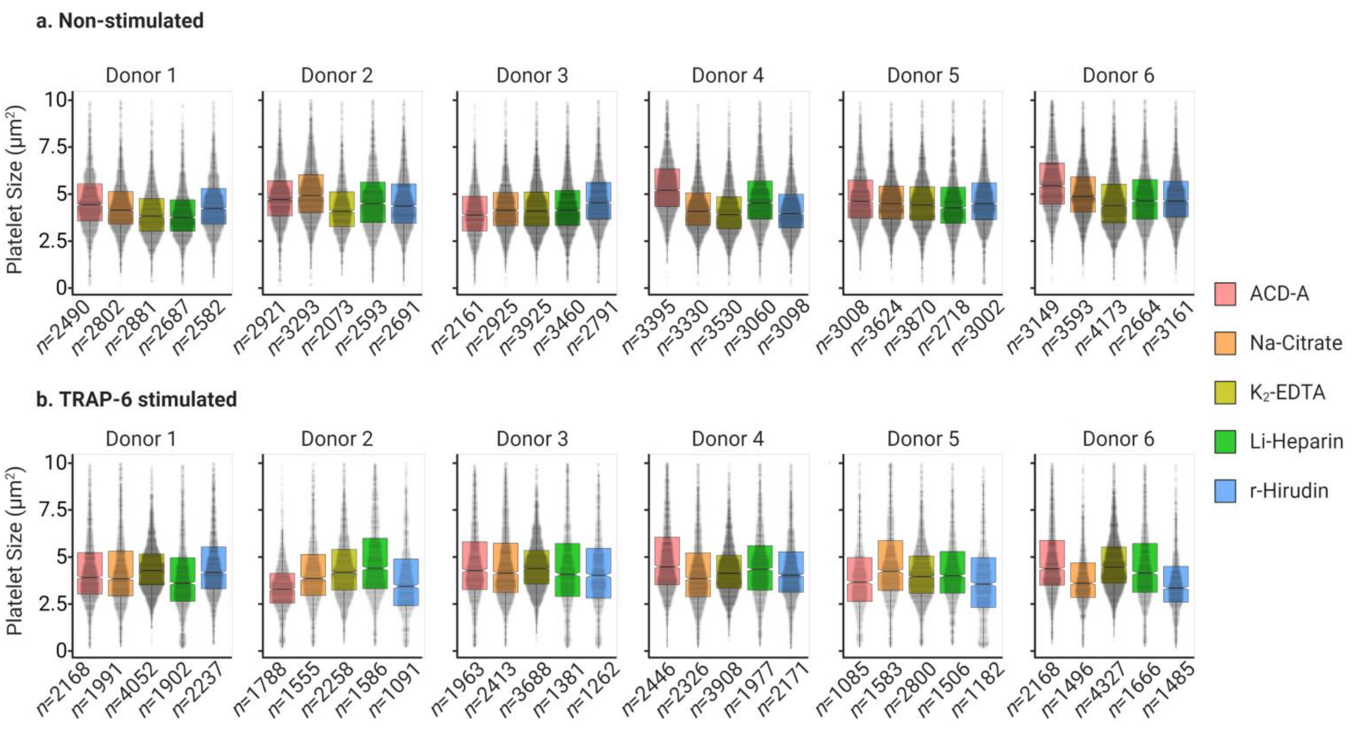


**Supplementary Fig. 5: Distribution of single platelet size in individual donors.** Violin plots overlaid with box plots of platelet size (µm$^2$) from six healthy donors before and after stimulation with PAR-1 agonist TRAP-6 peptide (20µM) for 10 minutes in the presence of PRP obtained from blood collected in different anticoagulants. n= number of single platelets analyzed for each per donor for each *ex vivo* anticoagulant.

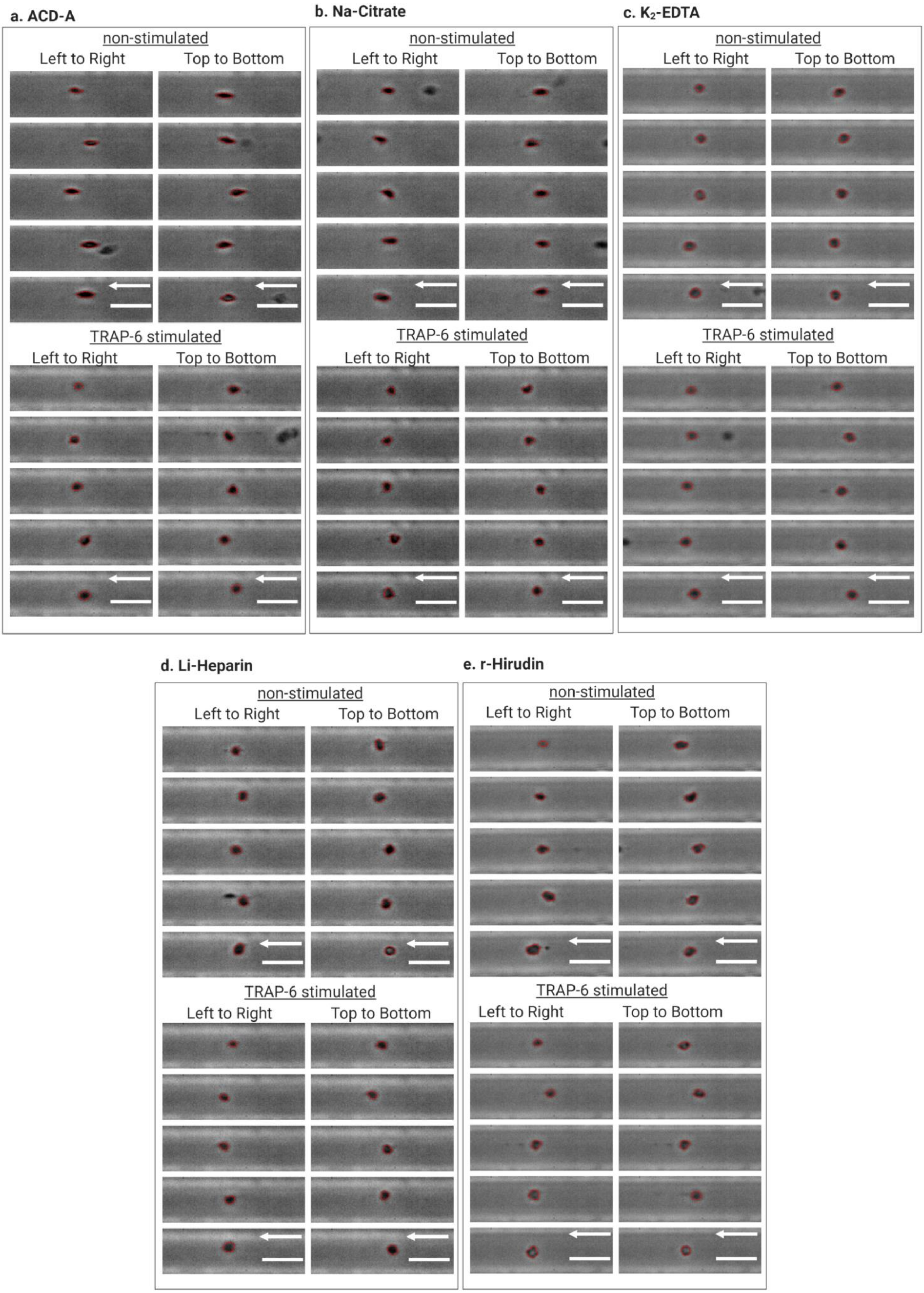

a. ACD-A
non-stimulated
Left to Right
Top to Bottom
TRAP-6 stimulated
Left to Right
Top to Bottom
b. Na-Citrate
non-stimulated
Left to Right
Top to Bottom
TRAP-6 stimulated
Left to Right
Top to Bottom
c. $K_2$-EDTA
non-stimulated
Left to Right
Top to Bottom
TRAP-6 stimulated
Left to Right
Top to Bottom
d. Li-Heparin
non-stimulated
Left to Right
Top to Bottom
TRAP-6 stimulated
Left to Right
Top to Bottom
e. r-Hirudin
non-stimulated
Left to Right
Top to Bottom
TRAP-6 stimulated
Left to Right
Top to Bottom

**Supplementary Fig. 6: Event images of platelet morphologies in the RT-FDC measurement channel.** Representative bright-field images of human platelets in different *ex vivo* anticoagulants **a.** ACD-A, **b.** Na-Citrate, **c.** K2-EDTA, **d.** Li-Heparin and **e.** r-Hirudin under non-stimulated (resting condition) and in the presence of platelet PAR-1 agonist TRAP-6. These event images were selected from RT-FDC scatter plots ( main text Figure 2-from left to right and top to bottom- 5 images each) of corresponding experimental conditions overlaid with the contour (red outline) in ShapeOut analysis software. Arrow indicates the direction of flow, and the scale bar corresponds to 10µm.

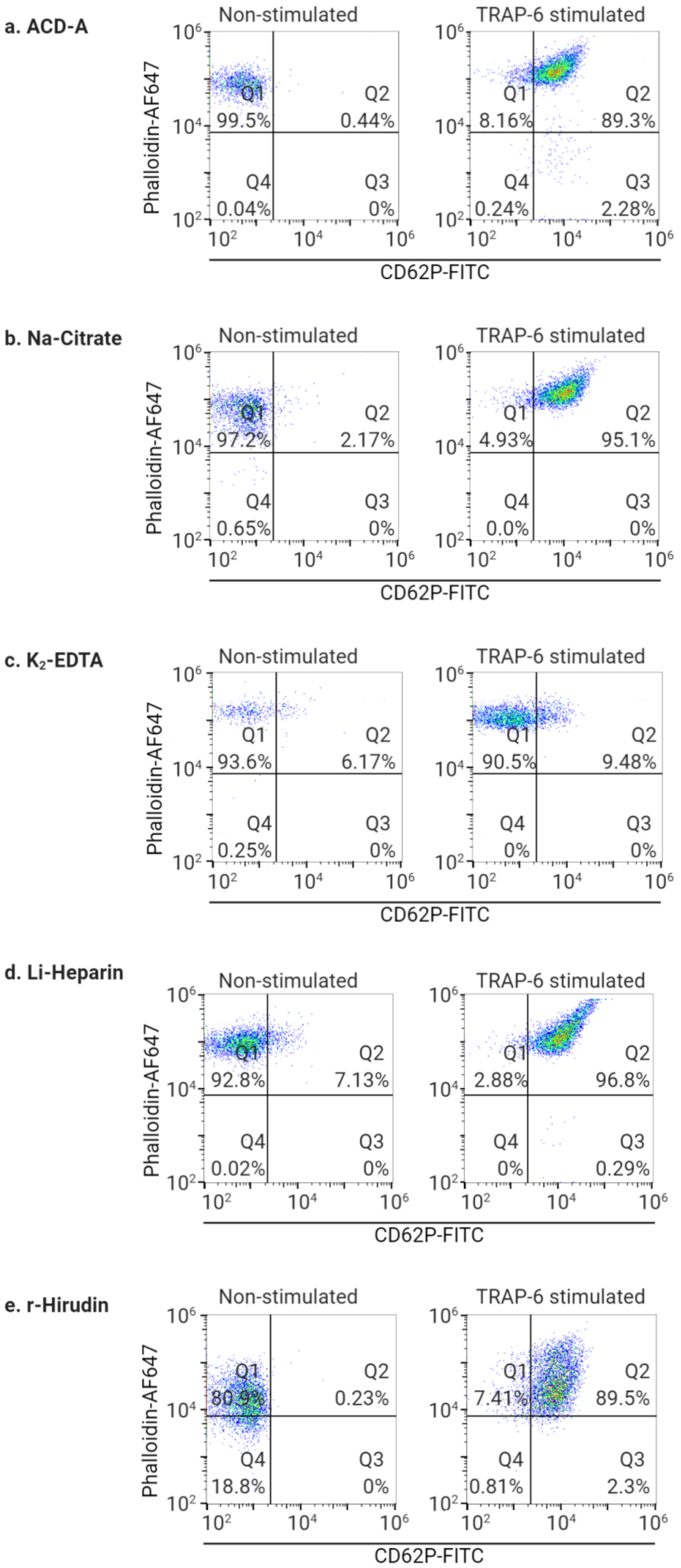

a. ACD-A
Non-stimulated
Q1 99.5%
Q2 0.44%
Q4 0.04%
Q3 0%
TRAP-6 stimulated
Q1 8.16%
Q2 89.3%
Q4 0.24%
Q3 2.28%
b. Na-Citrate
Non-stimulated
Q1 97.2%
Q2 2.17%
Q4 0.65%
Q3 0%
TRAP-6 stimulated
Q1 4.93%
Q2 95.1%
Q4 0.0%
Q3 0%
c. K2-EDTA
Non-stimulated
Q1 93.6%
Q2 6.17%
Q4 0.25%
Q3 0%
TRAP-6 stimulated
Q1 90.5%
Q2 9.48%
Q4 0%
Q3 0%
d. Li-Heparin
Non-stimulated
Q1 92.8%
Q2 7.13%
Q4 0.02%
Q3 0%
TRAP-6 stimulated
Q1 2.88%
Q2 96.8%
Q4 0%
Q3 0.29%
e. r-Hirudin
Non-stimulated
Q1 80.9%
Q2 0.23%
Q4 18.8%
Q3 0%
TRAP-6 stimulated
Q1 7.41%
Q2 89.5%
Q4 0.81%
Q3 2.3%
Phalloidin-AF647
CD62P-FITC

**Supplementary Fig. 7: Flow cytometric detection of platelet CD62P expression and F-actin content.** Representative flow cytometry bivariate density dot plots of non-stimulated and TRAP-6 stimulated platelets in PRP. Platelets are labeled for platelet specific activation marker CD62P (with anti-CD62P-FITC) and F-actin (with Phalloidin-AF647) in different *ex vivo* anticoagulants **a.** ACD-A, **b.** Na-Citrate, **c.** K2-EDTA, **d.** Li-Heparin and **e.** r-Hirudin. The quadrant regions show the percentage of platelets in each sub-population.

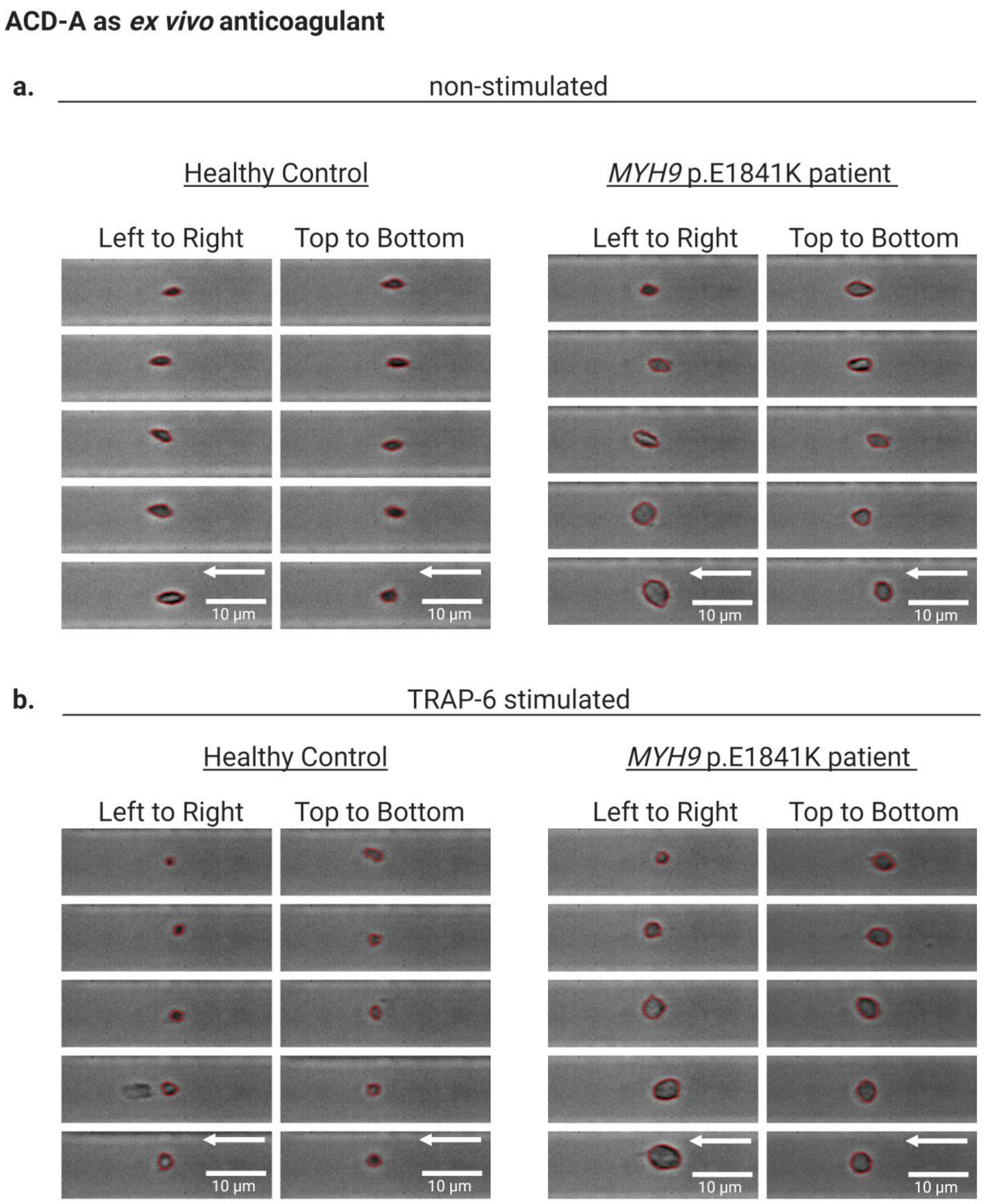


**Supplementary Fig.8: Event images with the contour of platelet morphologies from the RT-FDC measurement channel from healthy individual and MYH9 p.E1841K patient in ACD-A.** Comparison of **r**epresentative bright-field images of platelets in PRP prepared from ACD-A anticoagulated blood from a healthy individual and from a patient with MYH9 p.E1841K mutation under **a.** non-stimulated (resting condition) and **b.** in the presence of platelet PAR-1 agonist TRAP-6. These event images were selected from RT-FDC scatter plots ( main text Figure 7-from left to right and top to bottom- 5 images each) of corresponding experimental conditions overlaid with the contour (red outline) in ShapeOut analysis software. Arrow indicates the direction of flow and the scale bar corresponds to 10µm.

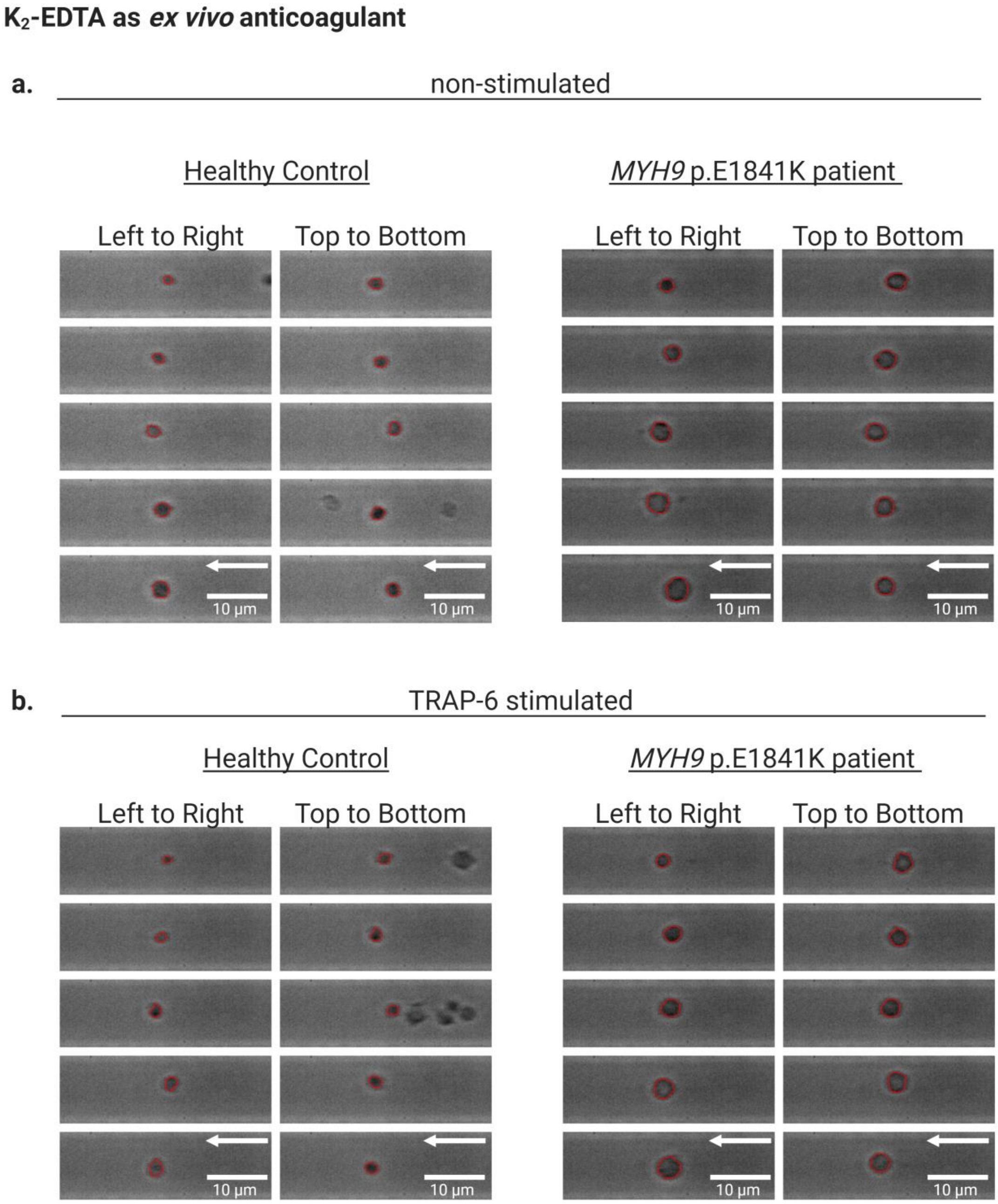


**Supplementary Fig. 9: Event images with the contour of platelet morphologies from the RT-FDC measurement channel from a healthy individual and MYH9 p.E1841K patient in ACD-A. in $K_2$-EDTA.** Comparison of **r**epresentative bright-field images of platelets in PRP prepared from $K_2$-EDTA anticoagulated blood from a healthy individual and from a patient with MYH9 p.E1841K mutation under **a.** non-stimulated (resting condition) and **b.** in the presence of platelet PAR-1 agonist TRAP-6. These event images were selected from RT-FDC scatter plots ( main text Figure 7-from left to right and top to bottom- 5 images each) of corresponding experimental conditions overlaid with the contour (red outline) in ShapeOut analysis software. Arrow indicates the direction of flow and the scale bar corresponds to 10µm.

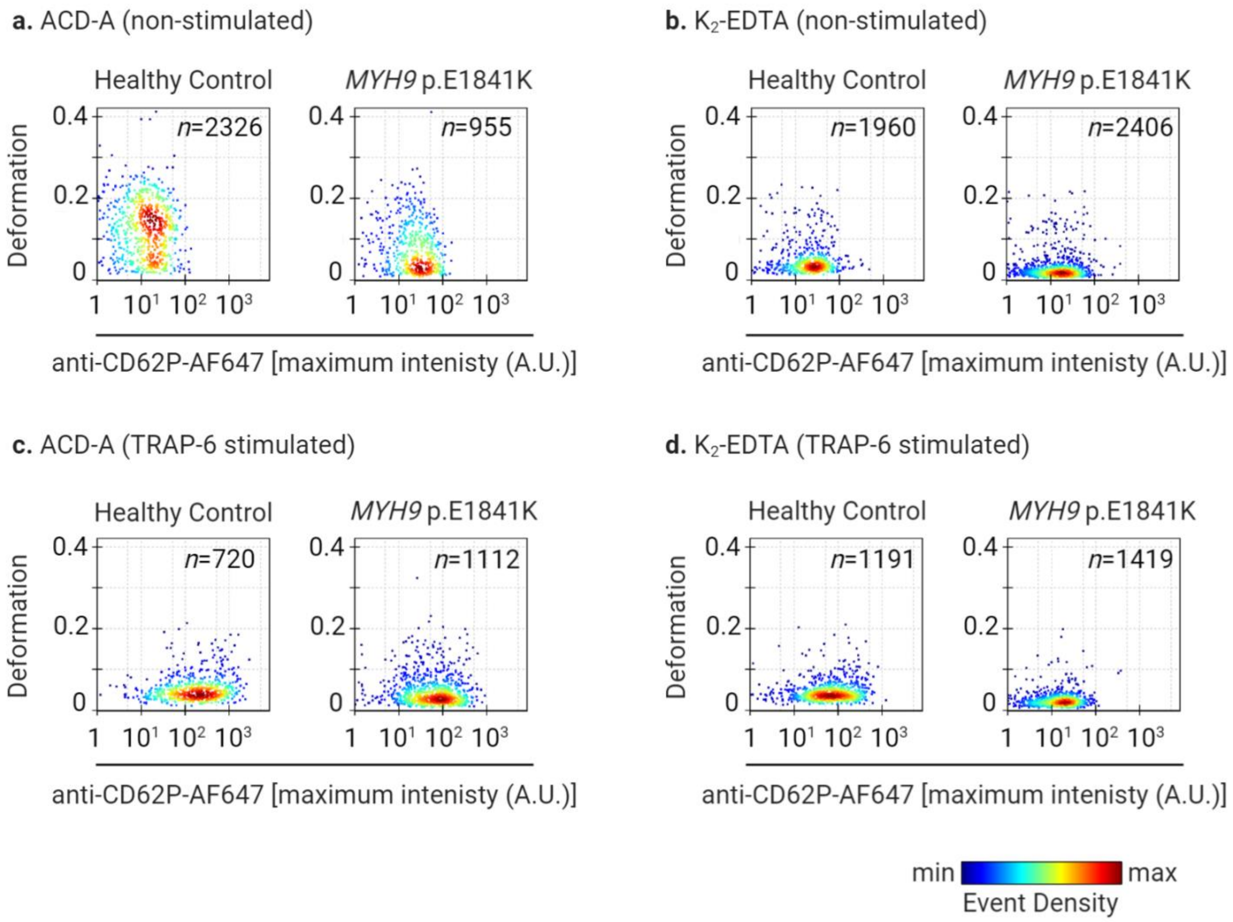


**Supplementary Fig.10: Platelet deformation and CD62P expression from patient with *MYH9* p.E1841K.** Comparison of scatter plots colored by Kernel Density Estimate (KDE) of RT-FDC data of deformation and CD62P expression (plotted on log10 scale of maximum intensity in arbitrary units (A.U.) of anti-CD62P-AlexaFluor647) on single platelets in PRP from a healthy individual and from a patient with MYH9 p.E1841K mutation under non-stimulated in **a.** ACD-A and **b**. K2-EDTA upon stimulation **c.** and **d.** with TRAP-6. Color coding of event density in scatter plots indicates a linear density scale from min (blue) to max (dark red).

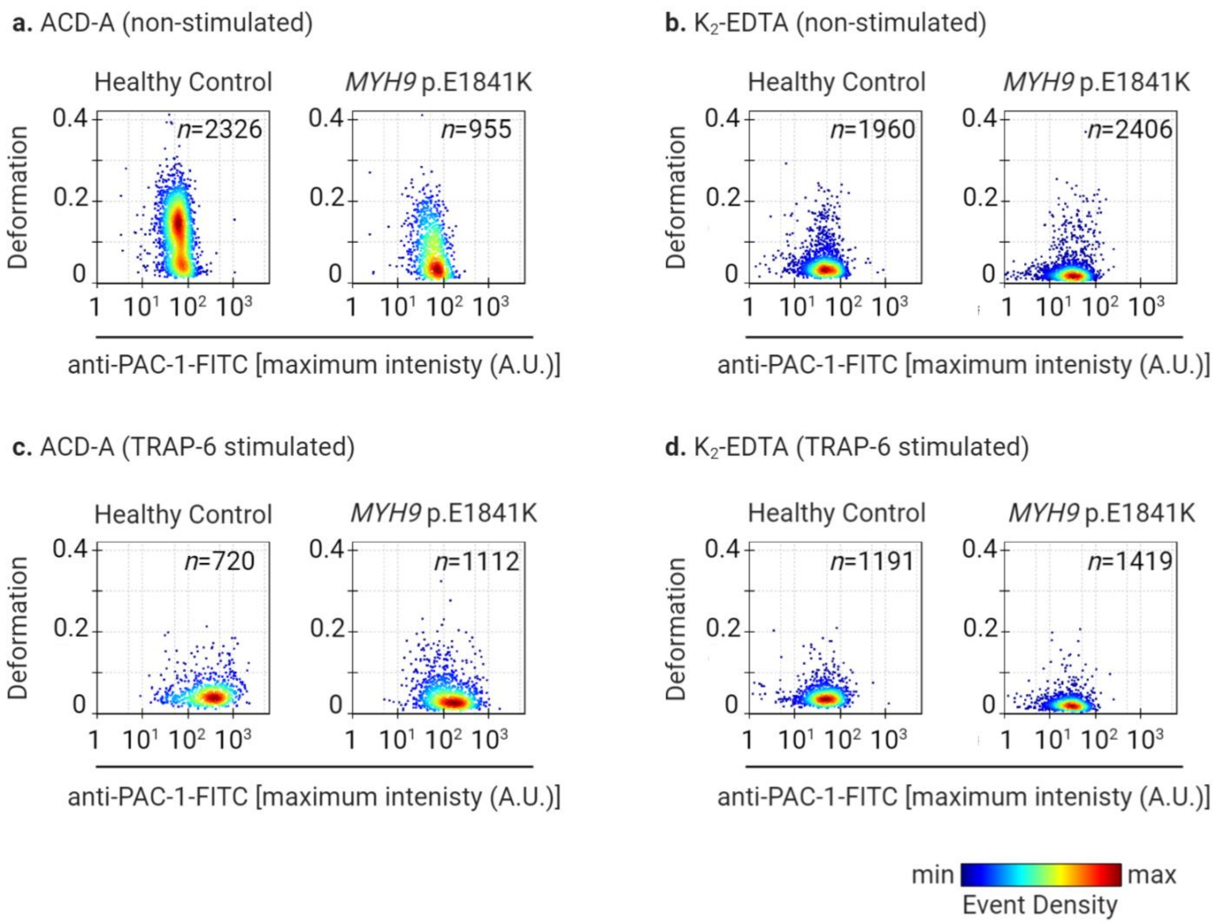


**Supplementary Fig. 11: Platelet deformation and platelet activation measured for binding of anti-PAC-1 antibody as a function of exposure of conformational epitope of the integrin αIIbβ3 from patient with *MYH9* p.E1841K.** Comparison of scatter plots colored by Kernel Density Estimate (KDE) of RT-FDC data of deformation and anti-PAC-1 binding (plotted on log10 scale of maximum intensity in arbitrary units (A.U.) of anti-PAC-1-FITC) on single platelets in PRP from a healthy individual and from a patient with MYH9 p.E1841K mutation under non-stimulated in **a.** ACD-A and **b**. K2-EDTA upon stimulation **c.** and **d.** with TRAP-6. Color coding of event density in scatter plots indicates a linear density scale from min (blue) to max (dark red).

**a. ACD-A**

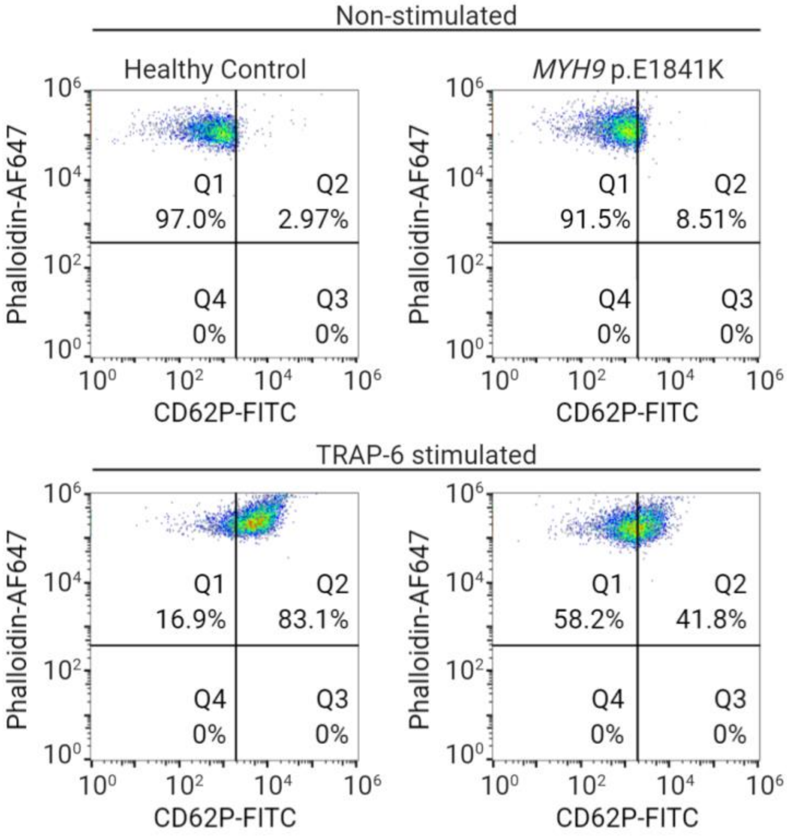


**b. $K_2$-EDTA**

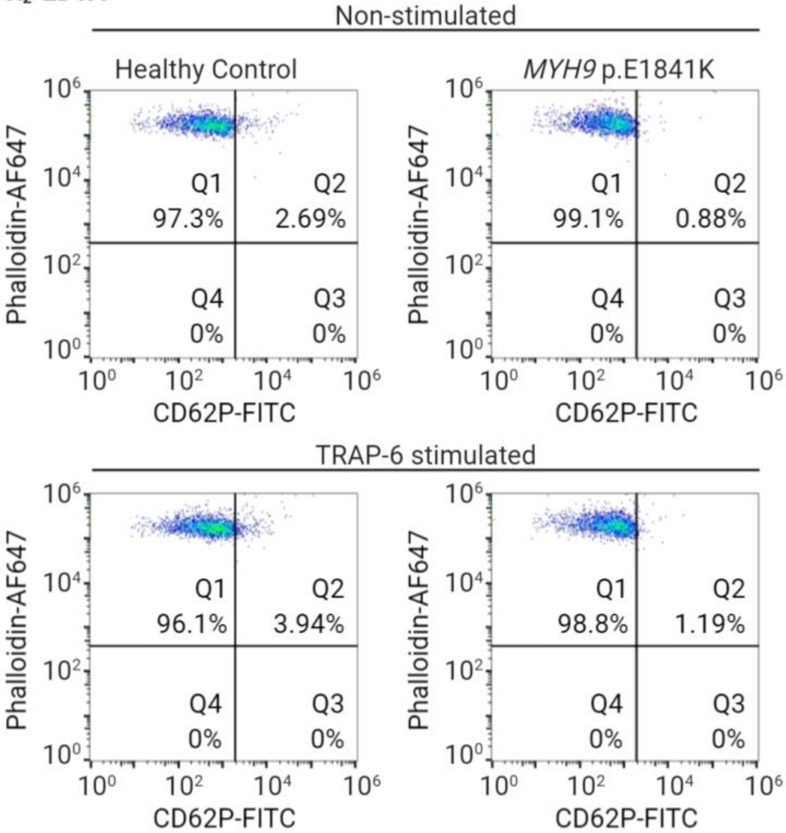

**Supplementary Fig.12: Flow cytometric detection of platelet CD62P expression and F-actin content in *MYH9* p.E1841K patient.** Flow cytometry bivariate density dot plots of non-stimulated and TRAP-6 stimulated platelets in PRP. Platelets are labeled for platelet specific activation marker CD62P (with anti-CD62P-FITC) and F-actin (with Phalloidin-AF647) in different *ex vivo* anticoagulants a. ACD-A and. b. $K_2$-EDTA.The quadrant regions show the percentage of platelets in each sub-population.
